# 3D observation of large-scale subcellular dynamics *in vivo* at the millisecond scale

**DOI:** 10.1101/672584

**Authors:** Jiamin Wu, Zhi Lu, Hui Qiao, Xu Zhang, Karl Zhanghao, Hao Xie, Tao Yan, Guoxun Zhang, Xiaoxu Li, Zheng Jiang, Xing Lin, Lu Fang, Bing Zhou, Jingtao Fan, Peng Xi, Qionghai Dai

**Affiliations:** Department of Automation, Tsinghua University, Beijing, 100084, China; Beijing Key Laboratory of Multi-dimension & Multi-scale Computational Photography (MMCP), School of Software, Tsinghua University, Beijing 100084, China; Department of Biomedical Engineering, College of Engineering, Peking University, Beijing 100871, China; Electrical and Computer Engineering Department, University of California, Los Angeles, CA 90095, USA; Tsinghua - Berkeley Shenzhen Institute (TBSI), Tsinghua University, Shenzhen 100084, China; State Key Laboratory of Membrane Biology, Tsinghua-Peking Center for Life Sciences, School of Life Sciences, Tsinghua University, Beijing 100084, China; Advanced Innovation Center for Big Data-Based Precision Medicine, School of Biological Science and Medical Engineering, Beihang University, Beijing 100191, China

**Author notes:** These authors contributed equally to this work (Jiamin Wu, Zhi Lu). Correspondence (Q. D.), (P. X.), (J. F.).

## Abstract

Observing large-scale three-dimensional (3D) subcellular dynamics *in vivo* at high spatiotemporal resolution has long been a pursuit for biology. However, both the signal-to-noise ratio and resolution degradation in multicellular organisms pose great challenges. Here, we propose a method, termed Digital Adaptive Optics Scanning Lightfield Mutual Iterative Tomography (DAOSLIMIT), featuring both 3D incoherent synthetic aperture and tiled wavefront correction in post-processing. We achieve aberration-free fluorescence imaging *in vivo* over a 150 × 150 × 16 μm^3^ field-of-view with the spatiotemporal resolution up to 250 nm laterally and 320 nm axially at 100 Hz, corresponding to a huge data throughput of over 15 Giga-voxels per second. Various fast subcellular processes are observed, including mitochondrial dynamics in cultured neurons, membrane dynamics in zebrafish embryos, and calcium propagation in cardiac cells, human cerebral organoids, and *Drosophila* larval neurons, enabling simultaneous *in vivo* studies of morphological and functional dynamics in 3D.

---

Cells in living organs compose an exquisite microscopic world in which the logistics, plasticity, interaction, and migration of multiple subcellular organelles can only be appreciated with high spatiotemporal resolution^1,2^. However, current microscopy techniques only image a certain plane at one time, with three-dimensional (3D) imaging obtained through movements of the focal plane relative to the specimen, such as confocal^3,4^, structured illumination^5,6^, light sheet microscopy^7–9^, etc. A few emerging imaging techniques^10^ aiming at simultaneous 3D imaging have been developed recently, such as light field microscopy (LFM)^11,12^, multi-focus microscopy^13,14^, etc. Although LFM has achieved great success in high-speed large-scale 3D calcium imaging^12,15,16^ with single-cell resolution, its resolution is typically limited to ~1 μm by the intrinsic trade-off between spatial and angular precision^17,18^, which is barely enough for subcellular structures. With holographic gratings, multi-focus images can be detected simultaneously at different areas on a large CCD camera^13,19^. However, they can only collect several discrete layers of the focal stack within a snapshot, and the total exposure time required for a high signal-to-noise ratio (SNR) is similar to that of axial-scanning methods due to the beam-splitting process. Considering the confined photon budget in fluorescence microscopy, the 3D imaging speed is crucially limited by the SNR of the imaging framework with the same exposure time^10^.

3D imaging of live cells requires not only high speed at high resolution but also faces an obstacle: the refractive index distribution changes along with the intracellular activity, creating significant aberrations by the crowded subcellular organelles. Such a common problem in multi-cellular organisms^2^ and biological tissues^20^ will reduce the resolution and SNR simultaneously, especially for objectives with high numerical aperture (NA)^21^. Adaptive optics (AO) is a technique commonly employed in telescopes against such distortions in which a microlens array and a 2D detector form a Shack-Hartmann wavefront sensor and an active phase modulator array is used for wavefront correction^21^. AO has successfully promoted the resolution of intravital optical microscopy to the diffraction limit^2^. However, one fundamental drawback for current AO is that, because the aberration changes over both time and locations, one can either use tiled AO for the entire imaging plane or compromise the correction mask by averaging phase distortions at different positions^22^. Applying a unique phase modulation to each pixel is too time-consuming, and sometimes it becomes impossible due to the lack of a guide star^21^. Although computational adaptive optics has achieved great success in optical coherence tomography^23,24^ and bright-field imaging, recently, it has been hard to apply in high-speed fluorescence imaging due to the incoherent property and low-photon condition.

To address all of these issues, we report digital adaptive optics scanning lightfield mutual iterative tomography (DAOSLIMIT), to realize aberration-free 3D fluorescence imaging at ultra-high spatiotemporal resolution across a large-scale volume. Based on LFM, instant 3D volumetric imaging can be acquired with high SNR in a tomographic way. The limited resolution of LFM can be compensated through the incoherent synthetic aperture of scanning LFM (sLFM). Moreover, we perform digital AO (DAO) with sLFM in post-processing, so that the phase distortion of every pixel can be corrected without compromising the data-acquisition speed, enabling real-time multi-site AO correction in fluorescence imaging, which was previously impossible. To avoid the loss of temporal resolution induced by scanning and further enhance the SNR for high-speed imaging, both time-loop and time-weighted algorithms are developed to fully exploit the redundancy of four-dimensional (4D) time-lapse video. The proposed DAOSLIMIT, to the best of our knowledge, provides orders-of-magnitude improvement in the spatiotemporal resolution for *in vivo* 3D fluorescence imaging with compact systems.

## Principle of DAOSLIMIT and experimental setup-up

Our DAOSLIMIT is built on a commercial epifluorescence microscope, with a microlens array inserted at the image plane for parallel acquisition of the low-resolution 4D phase space (Fig. 1a). Different sub-aperture components (labelled with different colours in Fig. 1a) at every local area are sampled by corresponding sensor pixels after each microlens. The microlens aperture is so small (approximately 5-6 times larger than the diffraction limit at the image plane), that it functions like the single-slit diffraction and produces frequency aliasing between different sub-aperture components^25^. The aliasing introduces coherence between the multiplexed phase space even for incoherent conditions, which is essential for the incoherent synthetic aperture. A two-dimensional galvo scanning system is utilized to shift the image plane by steps smaller than the pitch size of each microlens, which is equivalent to moving the microlens array relatively but in a much faster and more precise way. The scanning process introduces the spatial overlap of the local coherence measurements and increases the resolution of multiplexed phase-space measurements (Extended Data Fig. 1a, b). For illumination, a large volume within our 3D reconstruction range is excited simultaneously by a thick inclined illumination, which is effective for removing the out-of-range background fluorescence and preventing unnecessary photobleaching (Extended Data Fig. 1c). During data reconstruction, the sequentially scanned LFM images are first rearranged to get high-resolution multiplexed phase space (Supplementary Note 1 and Extended Data Fig. 2). Then, a mutual iterative tomography algorithm is developed to reconstruct the high-resolution 3D volume with the tiled wavefront correction for every pixel, which is critical for the inhomogeneous refractive index in multicellular organisms.

**Fig. 1.**
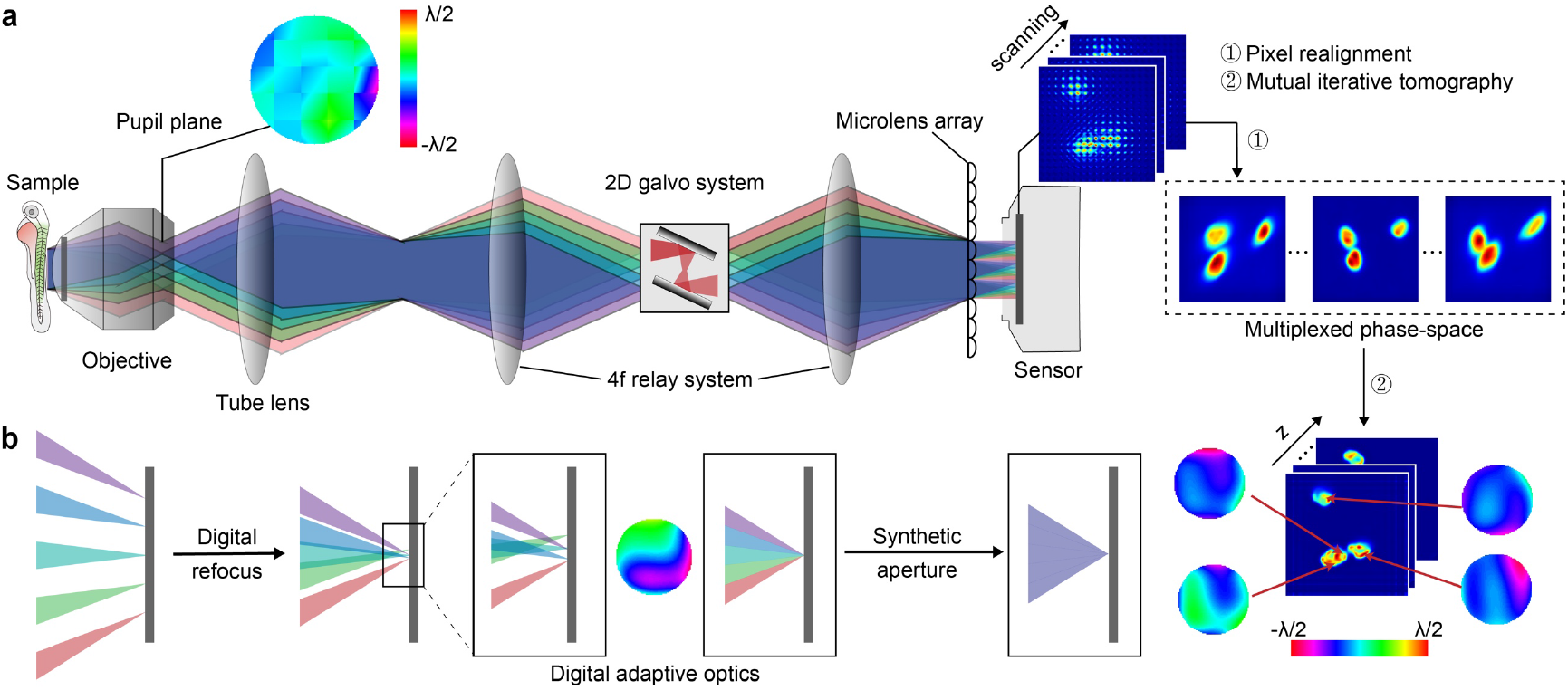
Schematic of digital adaptive optics scanning lightfield mutual iterative tomography (DAOSLIMIT). **a**, The system works in epi-fluorescence mode. A large volume within the 3D imaging range is excited simultaneously by a highly-inclined illumination and collected by the DAOSLIMIT. The inhomogeneous distribution of the refractive index in multicellular specimens produces strong spatially nonuniform aberrations at the back-pupil plane, which can be segmented for correction with adaptive optics. For illustration, light from different sub-apertures is labelled with different colours. A microlens array is inserted at the image plane for parallel acquisition of multiplexed phase-space measurements, whose resolution is further enhanced by the scanning process with a two-dimensional galvo scanning system. During reconstruction, we first realign the pixels from the raw data into the high-resolution multiplexed phase-space. Then, a mutual iterative tomography algorithm is employed to obtain the high-resolution volume with pixel-wise wavefront corrections. **b**, The multiplexed phase-space measurements can be synthesized for 3D reconstruction with the digital beam propagation. However, the sample-induced aberration will result in a distorted focus. We can digitally shift the sub-aperture point spread function (PSF), akin to applying a correction wavefront estimated during the volume reconstruction, to create a perfect focus. Both the spatial overlap induced by scanning and the frequency aliasing induced by the small aperture of each microlens facilitate the incoherent synthetic aperture during the 3D reconstruction, up to the diffraction limit of the whole objective’s NA.

The 4D phase-space measurements facilitate digital beam synthesis in both coherent ^26^ and partially coherent conditions^27^, providing sufficient information for 3D reconstruction (Fig. 1b, Supplementary Note 2, and Extended Data Fig. 3). However, the aberrations induced by the sample and optical system itself result in a distorted focus^21^ that was hardly recognized in previous LFM due to the reduced spatial resolution. To create a perfect focus, we digitally shift the sub-aperture point spread function (PSF), because a linear phase modulation at the segmented rear pupil function represents a specific lateral shift of the corresponding sub-aperture PSF (Supplementary Note 3). Such a process is analogous to AO via pupil segmentation by applying an estimated correction wavefront^20^. In addition, both the overlap in spatial domain induced by the scanning and the frequency aliasing in the angular domain facilitate the incoherent synthetic aperture during 3D reconstruction. As with the techniques sharing similar support constraints for recovery, such as ptychography^26,28^ and Wigner-distribution deconvolution^29^, we can achieve the diffraction limit of the whole objective NA with iterative updates of a consistent volume (Supplementary Note 4, and Extended Data Fig. 4).

## Results

### 3D synthetic aperture with digital adaptive optics for incoherent conditions

By measuring the local variance of coherence, phase-space imaging setups provide a computational method for 3D imaging, especially for partially coherent and incoherent conditions^27,30^. However, even with advanced algorithms^17,25,31^, the reconstructed lateral resolution without sacrificing the depth-of-field (DOF) is still limited to ~1 μm due to the failure of aperture synthesis in partially coherent and incoherent conditions such as fluorescence microscopy^10^. To mitigate the fundamental challenge, multiplexed phase-space measurements with coded aperture techniques have recently been employed ^32,33^, but the intensity modulation with a small transmission ratio leads to low light efficiency and loses the robustness and flexibility of segmented apertures. Previous methods utilizing scanning schemes^34,35^ enhance the resolution of perspective views to some extent, but they are still far from the diffraction limit of the whole objective’s NA in 3D reconstruction. With the high-resolution multiplexed phase-space measurements captured by DAOSLIMIT, we can apply the deconvolution algorithm for incoherent diffraction-limited 3D synthetic aperture, which is impossible for traditional LFM (Supplementary Note 2 and Extended Data Fig. 3).

Meanwhile, despite many investigations of the computational phase modulation capability with phase-space measurements, their applications are still limited in low-resolution extended depth of field^36^ and digital refocusing^11^. In particular, the unique advantage of phase-space measurements in wavefront correction has rarely been recognized, since the image degradation from the aberration is often concealed by the reduced resolution of traditional phase-space synthesis. However, with our diffraction-limited incoherent synthetic aperture capability, we fully exploit the power of computational phase modulation in post-processing, which is called digital adaptive optics (DAO). Without using a deformable mirror^20^ or spatial light modulator (SLM)^22^ as in traditional AO, we computationally apply multi-site aberration corrections to further enhance the resolution and SNR (Supplementary Note 3).

For high-speed 3D fluorescence imaging as shown in Fig. 2a, rather than the axial scanning by normal wide-field fluorescence microscopy (WFM), DAOSLIMIT captures the 3D signals along multiple elongated PSFs without any optical loss by lateral scanning (Fig. 2a). To realize diffraction-limited 3D incoherent synthetic aperture and DAO together, we develop an algorithm, termed mutual iterative tomography, that incorporates both the iterative wavefront estimation and volume reconstruction with digital correction together, by the alternating direction multipliers (ADMM) method^37^ (Supplementary Note 4 and Extended Data Fig. 2). Even with a larger input data size due to the higher resolution of phase-space measurements, we get similar convergence speed to the traditional 3D deconvolution algorithm but retrieve much more effective voxels (at least 30 times, Extended Data Fig. 5a, b).

**Fig. 2.**
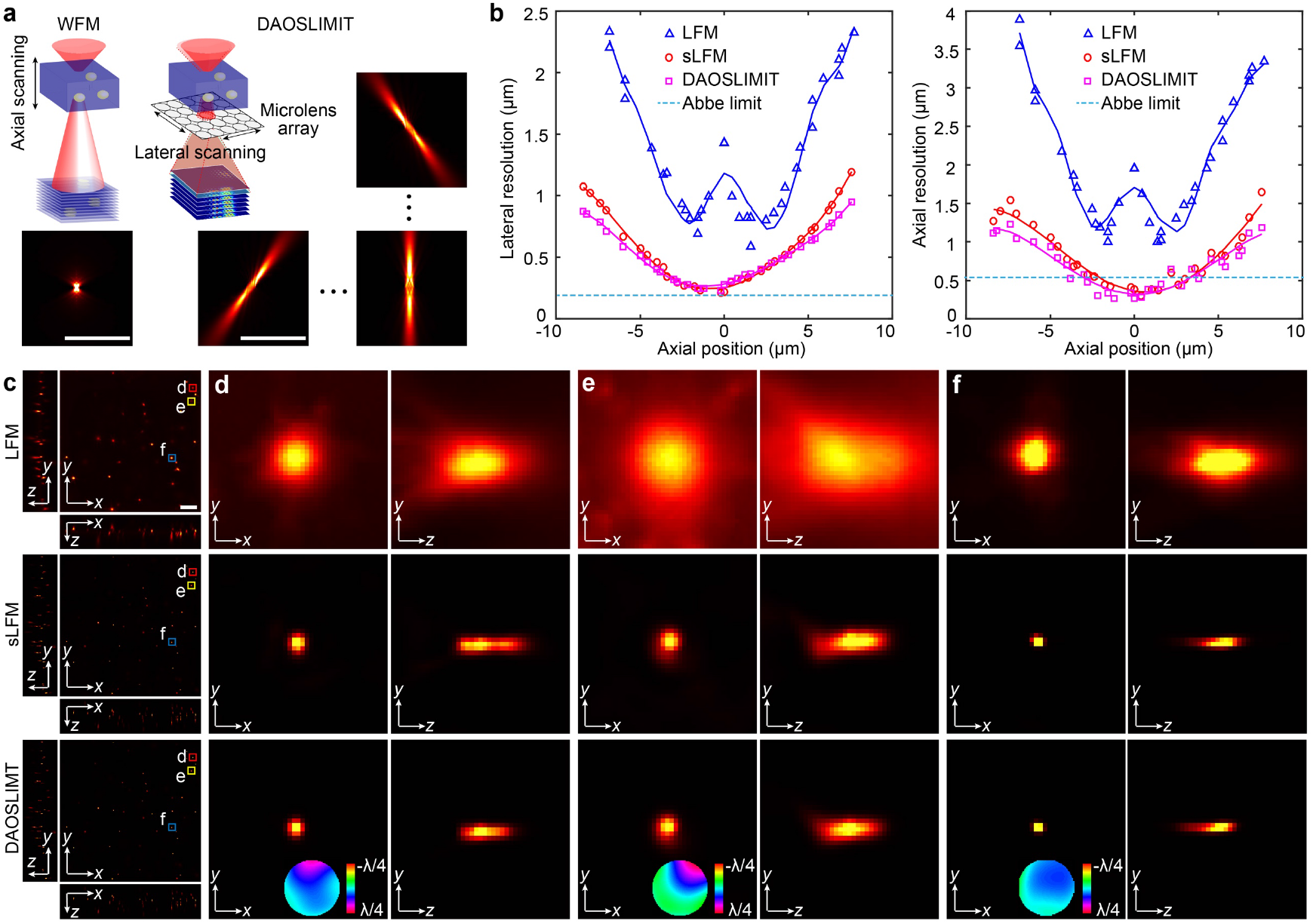
Resolution characterization in 3D fluorescence imaging. **a**, Different sampling strategies for wide-field fluorescence microscopy (WFM) and DAOSLIMIT. The PSFs of DAOSLIMIT for different spatial frequency components all have a larger depth of field than the WFM, illustrating the multiplexing capability in a tomographic way. Scale bar: 10 μm. **b**, We characterize the system resolution with 100-nm-diameter fluorescence beads distributed in 1% agarose. For beads at different axial positions, we calculated the lateral and axial resolution by measuring the full width at half-maximum (FWHM) of the lateral and axial profiles with Gaussian fit, respectively. The resolution of LFM is shown by the blue curves with triangles, whereas those of sLFM only and DAOSLIMIT are shown by the red curves with circles and the pink curves with rectangles, respectively. The Abbe limit is shown with the blue dashed line for comparison. **c**, Orthogonal maximum intensity projections (MIPs) of 100-nm-diameter fluorescence beads distributed in 1% agarose reconstructed by LFM, sLFM, and DAOSLIMIT, respectively. Scale bar: 10 μm. **d**–**f**, Zoom-in views of different fluorescence beads marked in c. The multi-site correction wavefronts are shown at the bottom. Scale bar: 1 μm.

To quantitatively evaluate the resolution after reconstruction, we imaged 100-nm-diameter fluorescence beads distributed in 1% agarose with a high-NA objective (63×/1.4-NA) by traditional LFM, sLFM without DAO and DAOSLIMIT. For beads at different axial positions, we calculated the resolution by measuring the full width at half-maximum (FWHMs) of the lateral and axial profiles after Gaussian fit (Fig. 2b). With approximately 92% spatial overlap, sLFM achieves around three-fold improvement on average in both lateral and axial resolution, compared with traditional LFM. With DAO, the resolution can be further enhanced by correcting the spatially nonuniform aberrations, which are hard to recognize in traditional LFM due to resolution loss (Fig. 2c-f). Traditional LFM has a significant resolution drop close to the native object plane, as a trade-off for a larger depth of field, which is hard to bypass due to the resolution limitation in multiplexed phase space^17^. In contrast, our DAOSLIMIT has a much smoother curve with an even larger depth of field. Both the lateral and axial resolutions reach the diffraction limit around the native object plane (250 nm and 320 nm, respectively) and maintain similar performance over a large axial distance (~10 μm). The slight super-resolution capability in the axial domain close to the focal plane is mainly due to the higher angular sensitivity and deconvolution process, which is identical to the previous study with even smaller angular resolution^38^. In addition, we tested the influence of different overlap ratios in the spatial domain and found that a similar performance can be achieved with only 67% spatial overlap, corresponding to 3×3 lateral shifts (Extended Data Fig. 6a, b), which is much smaller than the number of reconstructed axial planes (>100). Such data compression in a physics model reduces the sampling points required for 3D imaging, benefiting from the repeated usage of every pixel in phase space at different axial planes with the prior of linear beam propagation.

### Enhanced SNR for high-speed, high-resolution 3D fluorescence imaging

Rather than acquiring the focal stack in WFM with limited depth of field for each focal plane, DAOSLIMIT significantly improves the SNR by multiplexing different axial planes effectively along various sub-aperture PSFs, which stay focused within the extended depth of field^25^. To verify the SNR enhancement, we imaged HeLa cells labelled with GFP on the actin and DAPI on the nuclei, by DAOSLIMIT and WFM (Fig. 3). Every image of the WFM (first) and DAOSLIMIT (later) was captured sequentially with the same excitation power and exposure time for a fair comparison. A small region was cropped out with 4 cells for detailed analysis. Low excitation power (Supplementary Table 1) was utilized to create a low-SNR condition and avoid photobleaching. The orthogonal maximum intensity projections (MIPs) across 16-μm-thick slabs of the focal stack acquired by WFM, including 90 axial slices at 200-nm steps (corresponding to 4.5 s total exposure time), show strong background fluorescence and great shot noise with extremely low contrast in the *x-z* and *y-z* planes (Fig. 3a). Employing 3D Richardson-Lucy (RL) deconvolution^39^, we significantly enhance the contrast and SNR with the 3D wide-field PSF incorporating the photons from 90 images, which can be clearly observed from its Fourier transform (Fig. 3b). Fewer images used for WFM decrease the SNR and resolution, as shown in Extended Data Fig. 7a-d. Conventional LFM obtains the 3D information in a snapshot at the expense of spatial resolution (Fig. 3c), with some reconstruction artefacts close to the native objective plane^25^. Our DAOSLIMIT pushes the resolution to the diffraction limit with much higher SNR and contrast than other methods (Fig. 3d), and only 9 images are used here for reconstruction (10 times less than WFM). Similar subcellular structures can be observed in both Figs. 3b and d, while the DAOSLIMIT results show even better axial resolution with much less noise.

**Fig. 3.**
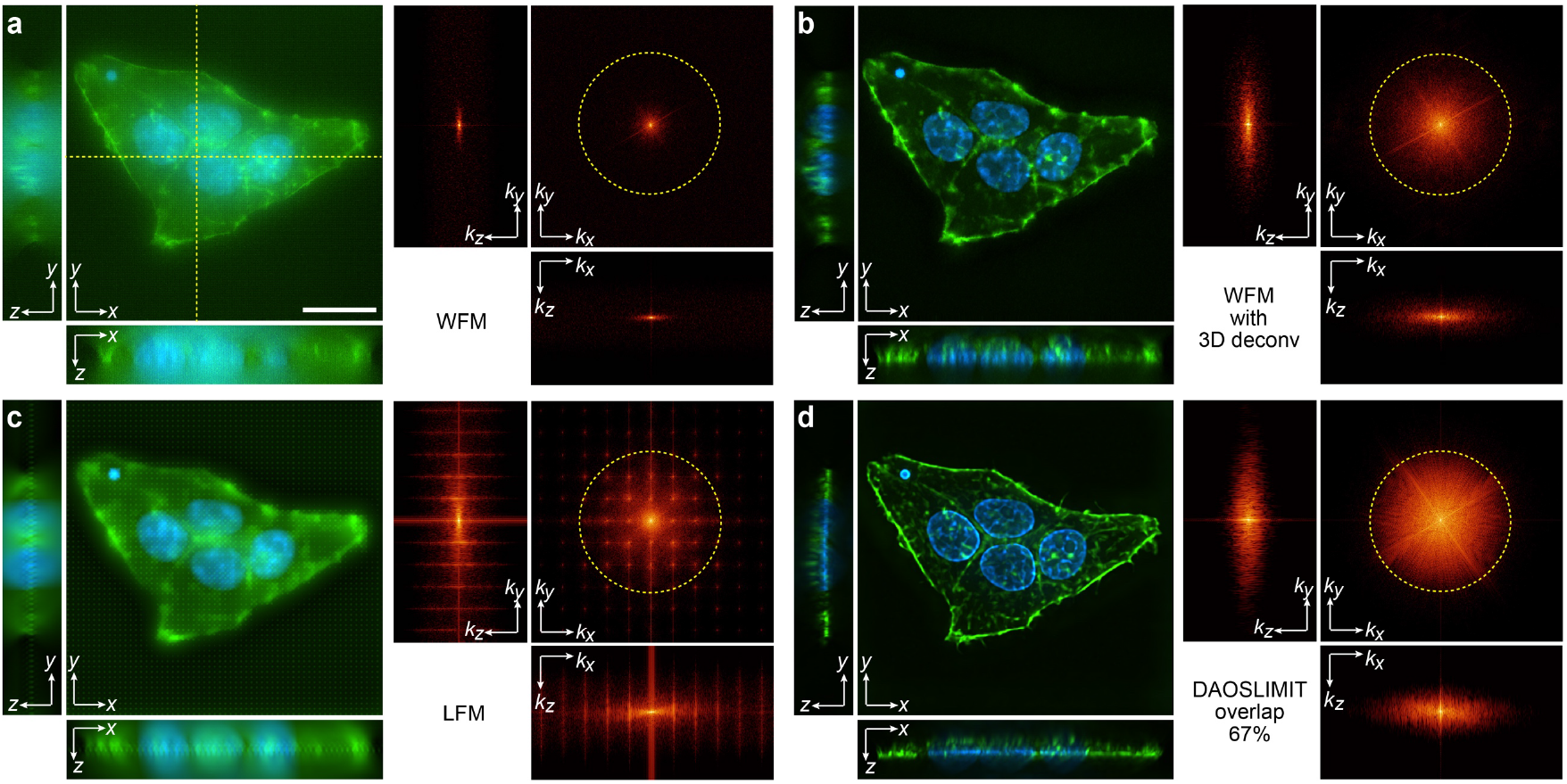
SNR and contrast improvements by DAOSLIMIT in 3D fluorescence imaging. **a**, Orthogonal MIPs from 16-μm-thick slabs of the focal stack captured by WFM, including 90 axial slices at 200-nm steps. The sample used here is a HeLa cell labelled with actin (green) and nuclei (blue). The Fourier transforms of the MIPs are shown on the right to indicate the degrees of information recovery in low-SNR conditions. The yellow dashed circle corresponds to the spatial frequency of the Abbe diffraction limit for comparison. **b**, Results reconstructed by applying 3D RL deconvolution to the focal stack in a, with enhanced SNR and contrast. **c**, Results obtained by traditional LFM with 3D deconvolution, showing much lower spatial resolution and reconstruction artefacts close to the native object plane. **d**, Results reconstructed by DAOSLIMIT with a 67% overlap ratio (corresponding to 3 × 3 lateral shifts), indicating higher SNR and resolution especially in the *x-z* and *y-z* planes (notably only 9 images are used for 3D reconstruction, 10 times less than WFM). Scale bar: 20 μm.

Such a huge SNR improvement brought by the better 3D sampling strategy is essential for high-speed 3D fluorescence imaging^10^. We also compared the results of DAOSLIMIT with different spatial overlap ratios (Extended Data Fig. 7e, f), illustrating that spatial overlap larger than 67% offers little resolution enhancement. The ratio threshold is consistent with our results in resolution characterization and is similar to other ptychographic methods for aperture synthesis^26^. In addition, we demonstrated several comparisons with both commercial spinning disk confocal microscopy and light-sheet microscopy. The results show that we achieve much higher SNR at similar speeds with low photobleaching, whereas other techniques degrade quickly without AO, especially for low-scattering samples (Extended Data Fig. 8a). For transparent areas (Extended Data Fig. 8b), DAOSLIMIT retains similar resolution and contrast to light-sheet microscopy at much higher speed with uniform performance across a large field of view (FOV). The orders-of-magnitude improvement means that we can either image at higher speed with the same duration, or image for much longer durations at the same speed as conventional methods with significantly less phototoxicity.

### Aberration-free 3D observation of mitochondrial dynamics at the millisecond scale

With superior spatiotemporal resolution and multi-site AO capability, we can observe large-scale organelle dynamics in 3D at the millisecond scale without aberration. For demonstration, we imaged 3D mitochondrial dynamics in dorsal root ganglion (DRG) neurons of mice across a large volume. The colour-coded MIPs of the reconstructed 3D volumes with and without DAO are shown in Figs. 4a and b, respectively. Due to the large size of the DRG neuron cell body and high NA of the objective, there are large aberrations and consequent image contrast degradation, especially for the mitochondria close to the nuclei, whose refractive index is much higher than the cytoplasm. In contrast, DAO achieves significant improvement in SNR and resolution with uniform performance across a large FOV by multi-site DAO (Fig. 4c). A typical region is selected to be viewed across 2-μm-thick slabs perpendicular to the detection axis (*xy*; left panels in Fig. 4d, e) or parallel (*yz*; right panels in Fig. 4d, e). For example, the white arrow in Fig. 4e indicates a ring structure that cannot be recognized in Fig. 4d due to aberration. Substantial improvement can be observed after tiled wavefront correction in both the *x-y* and *y-z* planes, with the correction wavefront marked on the upper-right corner. The cross-section profiles indicate a qualitative evaluation of diffraction-limited performance after DAO to distinguish two mitochondria with an extremely small interval. With the interleaved reconstruction, recently used in structured illumination microscopy (SIM)^40,41^, the imaging speed is approximately 17 volumes per second. Limited by the sensor pixel number (Andor Zyla 4.2 plus), each volume contains ~2048 × 2048 × 81 voxels, which is further cropped to the regions of interest for better visualization. As shown in Supplementary Video 1, large-scale various mitochondrial dynamics, such as fast movements (Fig. 4f) and fusion (Fig. 4g), can be clearly observed in 3D, with millisecond-scale temporal resolution and diffraction-limited spatial resolution. For a quantitative analysis, we applied a commercial tracking algorithm to the reconstructed 3D videos with and without DAO, and the 2D video of one axial plane in the middle with DAO (Fig. 4h). With the high-resolution high-speed 3D data reconstructed by DAOSLIMIT, more mitochondria can be tracked with longer tracking lengths. In addition, we observe a remarkable increase in average speed in 3D data, compared with the 2D data, because the projection from 3D to 2D reduces the amplitude of the speed vector. Thus, both the high-speed 3D imaging capability and real-time multi-site DAO are essential for robust and accurate tracking analysis of organelle dynamics.

**Fig. 4.**
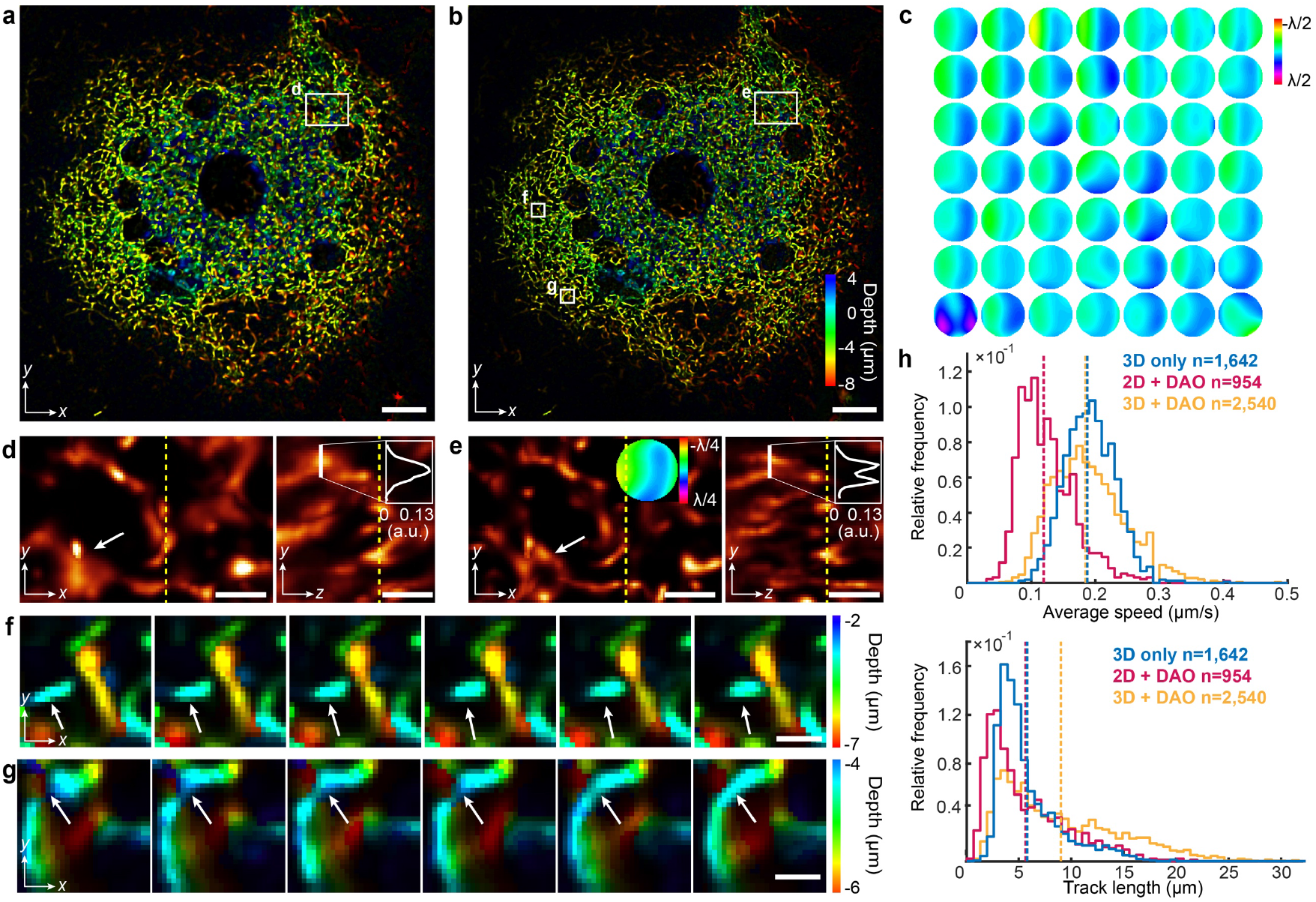
Millisecond-scale 3D fluorescence imaging of mitochondrial dynamics with DAO. **a**, Colour-coded MIP of the 3D volume reconstructed by sLFM without DAO. Different colours encode different depths. Scale bar: 10 μm. **b**, Colour-coded MIP of the 3D volume reconstructed by DAOSLIMIT. Scale bar: 10 μm. **c**, Different corrected wavefronts applied to different areas across the field of view in a. **d**, Orthogonal MIPs from 2-μm-thick slabs of the selected area in a, with a cross-section profile illustrating the resolution loss from aberration. The white arrow points to a structure blurred by the aberration. Scale bar: 2 μm. **e**, Orthogonal MIPs from 2-μm-thick slabs of the selected area in b, with a cross-section profile illustrating the diffraction-limited performance with DAO. The average correction wavefront of the selected region is shown in the inset at the unit of wavelength. The white arrow points to the ring structure resolved after DAO. Scale bar: 2 μm. **f**-**g**, Colour-coded MIPs of the selected areas in b at different time stamps marked at the bottom row. The arrow in **f** indicates a fast-moving mitochondrion in 3D, whereas the arrow in g indicates a mitochondrial fusion process happening in 3D. Scale bar: 1 μm. **h**, The distributions of the average speed and tracking length with the same tracking algorithm applied to 3D videos without DAO, 2D videos with DAO and 3D videos with DAO. More accurate speed can be estimated in 3D, whereas the SNR and resolution enhancements by DAO facilitate longer tracking duration and more mitochondria under tracking.

### *In vivo* imaging of vesicle and membrane dynamics during embryogenesis

To show the unprecedented imaging performance *in vivo*, we imaged various dynamics of vesicles and membranes in zebrafish embryos at the gastrulation stage. First, a comparison in Fig. 5a demonstrates that the resolution and contrast degradations of WFM in multicellular organisms can be successfully resolved by DAOSLIMIT with much fewer images required.

**Fig. 5.**
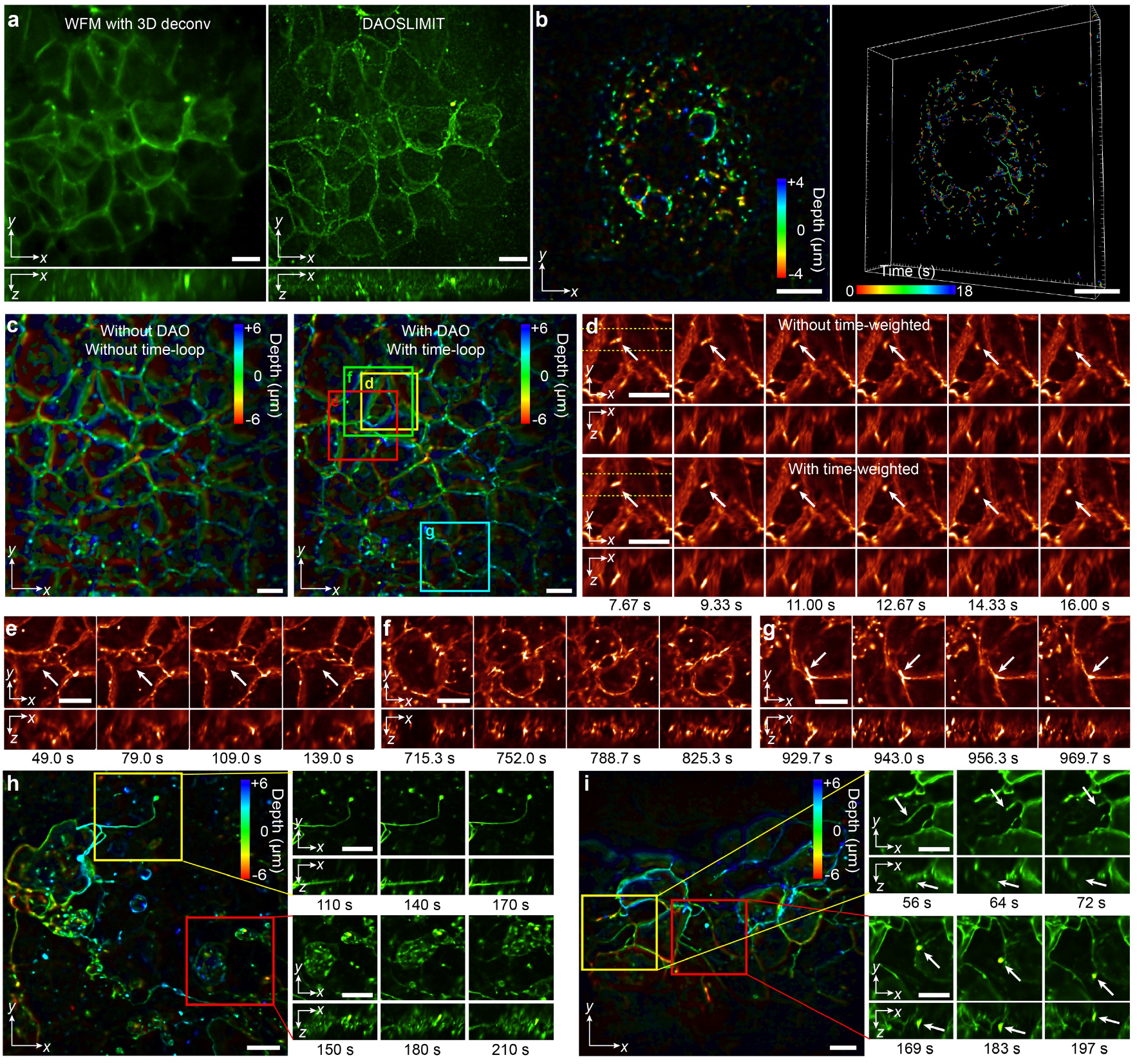
Vesicles and membrane dynamics in zebrafish embryos *in vivo*. **a**, Orthogonal MIPs of the 3D membrane structures reconstructed by WFM with 3D deconvolution (left panel) and DAOSLIMIT (right panel). **b**, 100 Hz 3D imaging of vesicle dynamics and potential vesicle trafficking in one zebrafish epithelial cell in the gastrulation stage (Supplementary Video 2). The colour-coded MIP of the volume at 0 s is shown in the left panel with different colours corresponding to different depths. The 3D tracking trajectories of every vesicle are shown in the right panel. **c**, The colour-coded MIP of the *in vivo* membrane dynamics reconstructed without DAO and time-loop algorithms are shown on the left, whereas that reconstructed with DAO, and time-loop algorithms is shown on the right, indicating improvement in both SNR and resolution (Supplementary Video 3). **d**, The effectiveness of the time-weighted algorithm on the compensation of the sacrificed temporal resolution resulting from lateral scanning (Supplementary Video 3). Orthogonal MIPs across 5-μm-thick slabs of the selected area in c are displayed with different time stamps marked at the bottom row. The results without the time-weighted algorithm are shown on the first row, and results with the time-weighted algorithm are shown on the second row, which eliminates the motion blur of a moving vesicle (white arrow) with the same spatial resolution. (**e-g**) Orthogonal MIPs of the selected areas in c with different time stamps marked at the bottom row (Supplementary Video 4). Dynamic filopodia retraction processes can be observed in e and g. A cell division process showed gradual membrane enrichment dynamics at the boundary of daughter cells in f. **h**–**i**, The colour-coded MIP of the 3D membrane dynamics *in vivo* at 10 Hz. Different zoom-ins with orthogonal MIPs in h show the 3D movements of a migrasome and a clear cell migration in 3D at high spatiotemporal resolution (Supplementary Video 6). High-speed fluctuations of the filopodia membrane can be observed in i, including filopodia retraction process and migrasome movements (Supplementary Video 7). Scale bar: 10 μm.

The temporal resolution sacrificed by lateral scanning can be further compensated by exploiting the low-rank sparsity in time-lapse (4D) data^40,41^. Unlike SIM with only part of the frequency domain captured for one shot, every frame of DAOSLIMIT has all the frequency information for the volume reconstruction with different periodic sparse sampling patterns in the spatial domain (Extended Data Fig. 3). In addition to such spectral-spatial data redundancy for synthesis, the reconstruction of the time-lapse data is regarded as a whole by utilizing the volume obtained from former time stamps as the initial sample prior for the current frame. To achieve uniform performance across the whole timeline, we updated the volume in a time-loop way with temporal continuity, similar to running a movie forward and backward again. Then, only 2 iterations are required for each volume and the SNR can be further enhanced by incorporating the information of all the frames. With interleaved reconstruction, we can push the speed of high-resolution large-scale 3D fluorescence imaging to the camera frame rates with extremely high SNR. A 100 Hz 3D imaging of vesicle dynamics and potential vesicle trafficking in one zebrafish epithelial cell at the gastrulation stage was demonstrated to show its superior performance (Fig. 5b and Supplementary Video 2). The high spatiotemporal resolution facilitates high-fidelity 3D tracking of large-scale vesicles in a single cell with a large variance in speed from 0.2 μm/s to 10 μm/s. Our DAO capability provides strong robustness to optical aberrations and keeps diffraction-limited performance in multicellular organisms, while the time-loop algorithm further enhances the SNR and contrast with reduced computational costs. Figure 5c (Supplementary Video 3) shows a typical example deep inside the zebrafish embryo. Different colours represent different axial positions in the colour-coded MIPs. The dynamic filopodia retraction processes can be observed in Figs. 5e, and g (Supplementary Video 4). A cell division process (Fig. 5f) happened nearby and showed gradual membrane enrichment dynamics at the boundary of daughter cells. However, for samples with high-speed movement or intensity changes, the interleaved reconstruction only brings motion blur (Fig. 5d) or reconstruction artefacts (Extended Data Fig. 9). To effectively improve the temporal resolution, we develop a time-weighted algorithm by applying different weights to the sequentially captured light field images in the pre-processing step for high-resolution phase space. Orthogonal MIPs across 5-μm-thick slabs of the selected area in Fig. 5c are displayed to show the effectiveness of the time-weighted algorithm. The motion blur and artefacts of a fast-moving vesicle (white arrow) can be eliminated, and the surrounding structures are consistent without any resolution reduction. We further tested the time-weighted algorithm by imaging the histone-labelled *C. elegans*, which was highly dynamic and had obvious motion artefacts. With our time-weighted algorithm, little artefacts can be observed with clear 3D structures of histone in the nucleus (Extended Data Fig. 9 and Supplementary Video 5), which paves the way to study large-scale genome structure dynamics *in vivo*.

With both the time-loop and time-weighted algorithms, the migrasome dynamics and the migration process of a cell in 3D can be observed clearly at 10 Hz in Fig. 5h (Supplementary Video 6). We can also capture high-speed fluctuations of filopodia membrane (Fig. 5i, Extended Data Fig. 10 and Supplementary Video 7), including both fast filopodia retraction processes and 3D migrasome movements (Supplementary Video 7). High-speed 3D tracking can then be employed to analyse the motion patterns. Such experiments provide the high-throughput 4D data required for investigating cell-cell communications at the organelle level.

### 3D calcium propagation at subcellular resolution both *in vitro* and *in vivo*

Observing 3D calcium propagation at sub-cellular resolution *in vivo* has long been a pursuit in neuroscience^42^. Our DAOSLIMIT pushes the spatiotemporal resolution in multicellular organisms to a new stage, with orders-of-magnitude improvement, which provides a way to investigate the subcellular calcium dynamics both *in vitro* and *in vivo*. With the superior performance, cultured cardiomyocytes labelled with Fluo-8 were imaged at 100 Hz to show the 3D calcium propagation within a single cell (Extended Data Fig. 11 and Supplementary Video 8). For more challenging cases with large aberrations, we imaged the spontaneous calcium propagation in a human 3D cerebral organoid^43^ infected with the GCamp6s indicator at 30 Hz (Fig. 6a and Supplementary Video 9). We can clearly visualize the 3D calcium wave evoked from the intersection of two neurons, based on the temporal-coded MIP for start time, with different colours representing the time points when the calcium response (ΔF/F_0_) reaches 10% of the maximum intensity (Fig. 6b). Unlike the calcium increase in one neuron, a small local calcium reduction occurring in an adjacent neuron can be readily observed (Fig. 6e). In addition, with high speed, we can resolve the difference in rise time when the calcium intensity increases from 20% to 80% of the maximum intensity along the dendrite (Fig. 6c). The orthogonal MIPs of the selected area in Fig. 6a at different time stamps are shown in Fig. 6d. The temporal traces at the labelled regions of interest (ROIs) illustrate the diversity of calcium dynamics within a single neuron (Fig. 6e).

**Fig. 6.**
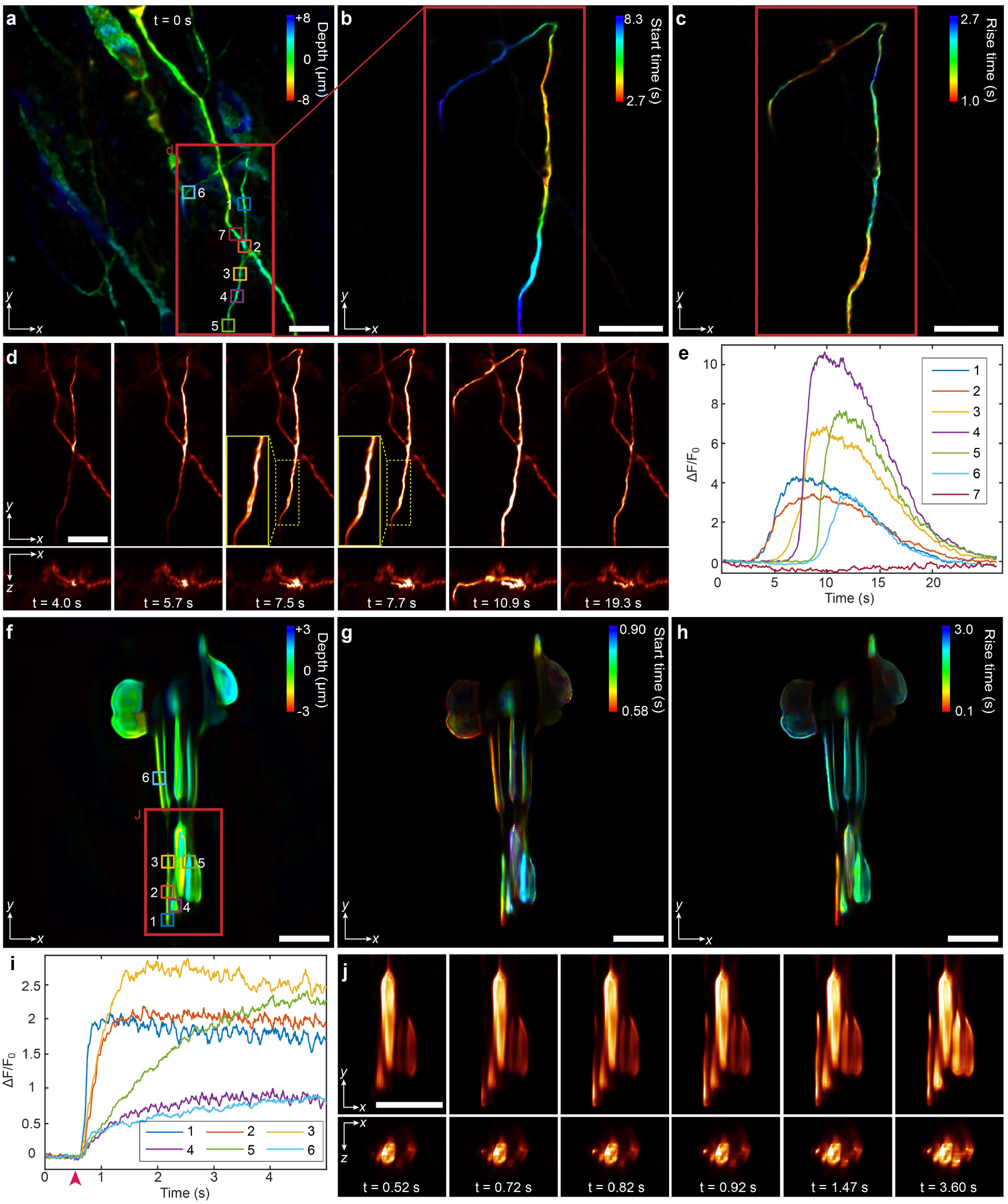
3D calcium propagation in human 3D cerebral organoids and *Drosophila* larval Cho neurons. **a**, Colour-coded MIP of the GCamp6s-labelled human 3D cerebral organoids obtained by DAOSLIMIT. Different colours represent different depths. **b**, Temporal-coded MIP of the selected area in a for the start time (the time point when the signal reaches 10% of the maximum intensity). Different colours correspond to the start points of the spontaneous calcium response, and the intensity represents the MIP of the temporal trace standard deviation for each voxel. Spontaneous 3D calcium propagation evoked from the intersection of two neurons can be clearly observed. **c**, Temporal-coded MIP of the selected area in a, illustrating the rise time (the time required to increase from 20% to 80% of the maximum intensity). Different colours correspond to the rise time of the spontaneous calcium response, and the intensity represents the MIP of the temporal trace standard deviation for each voxel. **d**, Orthogonal MIPs of the selected areas in a with different time stamps marked at the bottom row (Supplementary Video 9). The video was captured at 30 Hz. **e**, The temporal traces (ΔF/F_0_) of the ROIs labelled in a. **f**, Colour-coded MIP of the *Drosophila* larval Cho neurons with the jGCaMP7s indicator. Different colours represent different depths. **g,** Temporal-coded MIP for start time. **h**, Temporal-coded MIP for rise time. **i**, The temporal traces (ΔF/F_0_) of the ROIs labelled in f. The red arrow indicates the time point when we applied the 500 Hz sound stimulus. **j,** Orthogonal MIPs of the selected areas in f with different time stamps marked at the bottom row (Supplementary Video 10). The video was captured at 100 Hz. Scale bar: 10 μm.

For *in vivo* experiments, we conducted 3D calcium imaging of a cluster of five Cho neurons (lch5) in an awake *Drosophila* larva with sound stimulation^44^. Strong aberrations and fast movements of the larval body make it extremely difficult to record the neural activities at sub-cellular resolution in 3D. However, with DAOSLIMIT, we can easily obtain high-resolution 3D videos covering the whole Cho neuron at 100 Hz with sufficient SNR for detailed analysis (Fig. 6f and Supplementary Video 10). When we applied the 500 Hz sound stimulus (with the red arrow in Fig. 6i indicating the time point at which we started the stimulus), strong calcium responses showed up. The temporally coded MIP for start time illustrates that the speed of the calcium propagation varies a lot even on a same dendrite (Fig. 6g). In addition, a large variance of the rise time occurred in different parts of the lch5 neurons (Fig. 6h), which can also be observed in the temporal traces of the ROIs labelled in Fig. 6f. The zoomed-in orthogonal MIPs of the selected area in Fig. 6f at different time stamps are shown in Fig. 6j. With DAOSLMIT, more and more *in vivo* subcellular dynamics can be investigated in their native state across a large FOV with millisecond-scale temporal resolution.

## Discussion

We have demonstrated a highly efficient and compact microscopic imaging method to obtain aberration-free 3D fluorescence imaging at ultrahigh spatiotemporal resolution, termed digital adaptive optics scanning lightfield mutual iterative tomography (DAOSLIMIT). By multiplexing multiple spatial modes in a ptychographic way, we fundamentally solve the trade-off between the angular and spatial resolution in LFM. Such resolution promotion facilitates various computational imaging techniques such as 3D incoherent synthetic aperture and digital adaptive optics in fluorescence imaging, which, to the best of our knowledge, have never been realized before. With these unique capabilities, we achieve the first millisecond-scale aberration-free 3D fluorescence imaging at diffraction-limited resolution across a large volume *in vivo*, featuring over 15 Giga-voxels per second by data compression in the imaging side.

For WFM without aberration, every image has diffraction-limited resolution within the depth of focus. This traditional imaging scheme is extremely efficient for two-dimensional imaging with high resolution and contrast, although it does not work as well for 3D data sampling. Every slice in the focal stack has a small axial range with limited valid photons for 3D deconvolution. In contrast, sLFM keeps the PSF focused within the extended depth of field to avoid 3D information being flooded in the shot noise of the out-of-focus fluorescence, which is also a common benefit for PSF engineering in super-resolution techniques^45^. The photons from the 3D sample are then collected effectively in a tomographic way with robustness to scattering and aberrations benefitting from the low NA of the sub-aperture PSF. In addition, the scanning process, akin to ptychography, provides a strong constraint in the spatial domain for multiple state unmixing^46^. In addition to the efficient multiplexing mechanism, the repeated usage of high-resolution phase-space measurements in the aperture synthesis at different depths greatly reduces the required sampling points for a large volume, based on the assumption that the beams follow the propagation model within the depth of field^27^. The combination patterns are optimized by the DAO algorithm to get rid of the optical aberrations in a complicated environment and to reach the whole-NA resolution of the objective. The higher efficiency in both sampling and reconstruction, as well as the aberration-free capability with DAO significantly increase the SNR and the resolution of sLFM, compared with traditional WFM in multicellular organisms. Consequently, we believe such improvements in both SNR and spatiotemporal resolution will revolutionize the analysis and imaging techniques in various applications such as cell interactions in immune responses, cellular vesicular transport and high-speed 3D observation of calcium activity.

With DAOSLIMIT, we demonstrate fast aberration-free 3D volumetric imaging at diffraction-limited resolution, in a variety of biological specimens such as mitochondrial dynamics in live cells, vesicles and membrane dynamics in zebrafish embryos, and calcium propagation in human 3D cerebral organoids and *Drosophila* larval neurons. The capability of our DAOSLIMIT, working as a compact add-on to WFM, can be further exploited in multiple aspects. With diffraction-limited performance in multicellular organisms, structured illumination techniques^4,6^ can be incorporated to realize high-speed 3D super-resolution imaging *in vivo*. Moreover, different priors in spatial and temporal domains or the recently boomed deep learning techniques can be further associated with our technique for better performance. More generally, DAOSLIMIT can be extended to other aberration-sensitive imaging modalities such as gigapixel photography or X-ray computational tomography, to greatly improve their resolution, depth of field, and imaging speed simultaneously.

## Methods

### Experimental setup and imaging conditions

The DAOSLIMIT system works as an add-on to an inverted epifluorescence microscope (Zeiss, Observer Z1) equipped with a Metal Halide Lamp for fluorescence excitation. We used a 10×/0.45NA dry objective (Zeiss Apochromat) and a 10×/0.3NA phase objective (Zeiss Apochromat) for the USAF-1951 resolution chart, whereas all the biological experiments were performed with a 63×/1.4-NA oil objective (Zeiss Apochromat). A metal mask with a hollow ring is inserted at the pupil plane of the excitation path to reduce background fluorescence with a highly-inclined illumination^47^. The inner diameter and the outer diameter are chosen as 9 mm and 10mm, respectively, for the 63×/1.4-NA objective to excite a relatively-large volume covering the whole FOV (Extended Data Fig. 1). For the detection path, a 1.6× Optovar is chosen to get a further magnification of the image, which is then exported to the right-side image port of the microscope. We use a two-dimensional galvo-scanning system with a relay system between them to scan the image plane at both high speed and high precision. A microlens array with 100-μm pitch size and 2.1-mm effective focal length is then inserted at the conjugate image plane. Its NA (F-# 21) is slightly larger than the NA of the image plane to prevent spectrum leakage^11^. Another 4f system relays the back focal plane of the microlens array to the sCMOS camera (Andor Zyla 4.2 plus, 2,048 × 2,048 pixels) with a magnification of 0.845, so that each microlens covers 13 × 13 sensor pixels, corresponding to 1.43 μm × 1.43 μm area at the sample plane for the 63× objective. All the relay systems are custom-designed with off-the-shelf lenses to achieve diffraction-limited performance. Detailed imaging and reconstruction conditions for all fluorescence experiments in the paper, including excitation power, exposure time, frame rate, voxel sizes, fluorophores, proteins, and filter sets, are illustrated in Supplementary Table 1.

### Mutual iterative tomography with DAO

The volume reconstruction process can be viewed as an inverse tomographic problem along different angular PSFs^48^. Different from previous 3D RL deconvolution algorithm for LFM^25^, we model the problem in the 4D multiplexed phase space with details given in Supplementary Note 1 and Extended Data Fig. 2. The 4D PSF has several advantages over traditional 3D PSF including the spatial-invariance property and smooth distributions, which make the deconvolution process more efficient and robust.

The pipeline of the mutual iterative tomography algorithm is shown in Extended Data Fig. 4. For all the raw data recorded by the sensor, we first cropped the images to regions of interest (ROIs) with matched edges with the microlens. Then a pre-processing process was applied to realign the pixels in 4D multiplexed phase space. Pixels located at the same position relative to the centre of each microlens, in different sequentially-scanned images, were aligned together as a specific spatial frequency component. Before iterative updates, the realigned multiplexed phase-space data were up-sampled with a cubic interpolation as the input for reconstruction. To solve the optimization problem of 3D reconstruction with aberration estimation, we adopted the ADMM method^37^. Details of the algorithm are described in Supplementary Note 4.

### Time-lapse video reconstruction with time-loop and time-weighted algorithms

For time-lapse video, we recognized the reconstruction of the 4D information (the time-lapse 3D video) as a whole to make full use of the temporal continuity for SNR enhancement. An interleaved reconstruction process^40^ was applied to retrieve the temporal resolution sacrificed by the periodically scanning process (e.g. 3 × 3 lateral shifts by galvo). And we replaced the uniform initial value with the average of the uniform initial value and the reconstructed result of the previous frame, which could greatly accelerate the convergence. To get rid of the nonuniform performance with the increase of the frame number, we first updated the volumes with the evolution over time, and kept updating from the last frame to the first frame in the inverse direction, as a movie played back. We call such process the time-loop algorithm, which incorporates all the video information for every volume update and greatly improves the SNR due to the large redundancy in the 4D video. With only two iterations required for each volume at one time stamp, we can obtain the continuous 4D data at extremely high spatiotemporal resolution (Fig. 5c and Supplementary Video 3).

For samples with high-speed movements or flashing, we applied a time-weighted algorithm for pre-processing during the pixel realignment to alleviate motion blur (Fig. 5d and Supplementary Video 3) and artifacts (Extended Data Fig. 9 and Supplementary Video 5). For example, if we use 9 images to reconstruct a volume with 67% spatial overlap, images close to the middle time stamp will have higher weights during the up-sampling interpolation process. The time-weighted phase-space were then used as input for time-loop reconstructions described before, which can successfully reduce the motion blur as shown in Fig. 5d, indicating an effective improvement in the temporal resolution after interleaved reconstruction.

### Cell culture and slice preparation

Animal materials were collected in accordance with a protocol approved by the Institutional Animal Care and Use Committee of Tsinghua University.

HeLa cells were maintained in DMEM medium supplemented with 10% FBS, 1% Pen-Strep antibiotics and 1% NEAA (all from GIBCO) at 37 °C in a 5% CO_2_ incubator. 2 days before imaging, HeLa cells were seeded on the #1.5 Φ12 mm round cover glass (Fisher Scientific) at a density of 4000 cells per coverslip, then transfected by Cell Light™ GFP-actin reagents (Invitrogen). Upon imaging, cells were fixed in 4% paraformaldehyde (Sigma), washed by PBS and mounted by AntiFade mounting medium (containing 2 μg/mL DAPI, Invitrogen).

The cardiomyocytes were isolated from neonatal rats, and all the procedures were reviewed and approved by the Institutional Animal Care and Use Committee office of Tsinghua University. Briefly, rat ventricles were freshly isolated and minced into small pieces (0.5-1 mm^3^), washed with cold HBSS and digested in 200 U/mL Collagenase II (Worthington Biochemical Corporation) and 0.1% Trypsin (GIBCO) at 37 °C for 1h on the cell rotor, then neutralized by adding same volume of DMEM medium supplemented with 10% FBS (GIBCO). After filtered by 40 μm cell strainer (Falcon), cells were cultured in full DMEM medium for 2 h to remove the fibroblast and endothelial cells. The supernatant was centrifuged and cardiomyocytes were seeded on the #1.5 Φ12 mm round cover glass (Fisher Scientific) at the density of 6000 cells per coverslip, maintained at 37 °C in a 5% CO_2_ incubator. After 4 days, cells were washed by HBSS, stained with 2 μM Fluo-8 AM (AAT Bioquest) for 30 min, washed and imaged in HBSS at 37 °C.

For high-speed 3D imaging of mitochondrial dynamics, dorsal root ganglion neurons were trimmed and isolated from postnatal 8-10 day SD rat spinal cords in HBSS (Invitrogen), and digested in 2.5 U/ml dispase II (Roche) and 200 U/ml collagenase (Worthington Biochemical Corporation) at 37 °C for 30 min and then shaking on the cell rotor at 30 °C for 30 min. After a brief spin down, dissociate neurons were collected through 70-μm cell strainer (Falcon). For mitochondria labeling, 5×10^4^−1×10^5^ fresh isolated DRG neurons were transfected with 0.5 μg Mito-GFP for Nucleofector (Lonza) SCN Basic Neuro Program 6 with Amaxa Basic Neuron SCN nucleofector Kit, then seeded on 25 μg/ml Poly-ornithine (Sigma-Aldrich) and 5 μg/ml Laminin (Roche) pre-coated #1.5 Φ12 mm round cover glass (Fisher Scientific) at a density of 6000 cells per coverslip. After electroporation, neurons were maintained in Neurobasal A medium with 2% B27 supplements, 2mM GlutaMAX and 2% FBS (Invitrogen) for 48 h or 72 h at 37 °C in a 5% CO_2_ incubator before imaging.

### C. elegans experiments

The H2B::GFP strain AZ212 was kindly provided by Prof. Guangshuo Ou (Tsinghua University) as a gift and maintained on nematode growth medium (NGM) plates seeded with E. coli OP50 at 20°C following standard protocols. Live worm was imaged according to the previously described protocol^49^. Briefly, L1 larval worms were anesthetized with 1 mg/ml levamisole in M9 buffer and mounted on 3% agarose pads at 20°C. After immobilization, worms were imaged subsequently with our system.

### Zebrafish embryos experiments

Wild type (Tuebingen strain) zebrafish were used in this study with ethical approval from the Animal Care and Use Committee of Tsinghua University. The zebrafish embryos were injected with 300pg *tspan4a-EGFP* mRNA (synthesized *in vitro* with mMessage mMachine T7 kit (AM1344, Ambion) in one cell at the 16-cell stage (1.5hpf). At 70% epiboly stage (8hpf), injected embryos were embedded in 1% low-melting-point agarose in glass bottom dishes (D35-14-0-N, In Vitro Scientific) for live imaging. Fertilized zebrafish embryos were kept at 28.5°C in Holtfreter’s solution (NaCl 59 mM, KCl 0.67 mM, CaCl_2_ 0.76 mM, NaHCO_3_ 2.4 mM).

For the preparation of the fixed Zebrafish zebrafish gastrula used in Extended Data Fig. 8, 16-cell stage embryos were injected with about 300pg tspan4a-GFP mRNA in one cell. When embryos reached 70%-80% epiboly stage (7-8 hours post fertilization), we removed the chorion. −20 °C methanol was used to fix and dehydrate embryos overnight. The embryos were rehydrated by graded 0.5% PBST (Triton-X 100 Solarbio 524A0513) and then incubated in block solution (1% BSA, 0.3M glycine and 10% goat serum in 0.5% PBST) for 1 hour at room temperature. Embryos were then incubated overnight with primary antibody (chick anti-GFP (Abcam, ab13970)), which is diluted 1:50 in block solution at 4 °C. To remove the primary antibody, embryos were then washed in 0.5% PBST for 3×1 hour at room temperature and incubated in secondary antibody (anti-chick 488 (Abcam, ab150169)) diluted 1:200 in block solution overnight at 4 °C. To remove the secondary antibody, embryos were washed with 0.5% PBST for 3×1 hour at room temperature. Finally, the embryos were mounted in 1% low-melting-point agarose for imaging by DAOSLIMIT and the spinning-disk confocal microscope, respectively. The embryos were mounted directly for imaging by the light-sheet microscope.

### Human 3D cerebral organoids experiment

Human induced pluripotent stem cells (hiPSCs) DYR0100 were kindly provided by Stem Cell Bank, Chinese Academy of Sciences. Neural progenitor cells (NPCs) were differentiated from DYR0100 by Hopstem Bioengineering Ltd. Co following the differentiation method described previously^43^. Human 3D cerebral organoids were generated and maintained in human 3D cerebral organoids medium (Cat # HopCell-3DM-001, Hopstem Bioengineering) by Hopstem Bioengineering according to the manufacturer's protocols. For calcium imaging, human 3D cerebral organoids were infected with lentivirus carrying rLV-EF1α-GCamp6s-WPRE (provided by BrainVTA) and retuned to 37°C CO_2_ incubator for 1 week. After that, cerebral organoids were transferred to imaging chamber in calcium imaging buffer (150mM NaCl, 4mM KCl, 10mM HEPES, 10mM Glucose).

### *Drosophila* larvae experiments

The ChAT-Gal4 line (56500), UAS-TdTom (36328) and 20XUAS-IVS-jGCaMP7s line (79032) were obtained from the Bloomington Stock Center. Animals were raised at 25 °C in an incubator with a 12-h light/dark cycle and humidity control at 70%. *In vivo* calcium imaging of larval Cho neurons was performed with third instar larvae according to previously described protocols^44^. During imaging of DAOSLIMIT, a freely moving larva was pressed between two coverslips with a drop of distilled water to reduce its movement. The calcium indicator jGCaMP7s was used to measure the calcium signal.

### Data analysis

All the data analyses were done with customized Matlab (Mathworks, Matlab 2018) programs, Imaris 9.3 (Oxford Instruments) and Huygens. All the tracking results were done by Imaris with the same parameters for fair comparison. The 3D deconvolutions of the commercial light-sheet microscope and the spinning disk confocal microscope were done with Huygens. The temporal traces of calcium response are calculated by ΔF/F_0_= (F− F_0_)/F_0_, where F_0_ is the mean fluorescence intensity of the first 20 volumes. For all the colour-coded MIPs, we mapped the maximum intensity and its axial position to the value and hue in HSV colour space, respectively. For the temporal-coded MIPs, we firstly mapped the start time or rise time of each voxel to hue in HSV colour space, and then used the standard deviation of each voxel as the intensity for MIP calculation.

## Data availability

All relevant data are available from the corresponding authors upon reasonable request.

## Code availability

The codes for the reconstruction and analyses described in this paper are available from the corresponding author upon reasonable request.

## Acknowledgements

We thank Guangshuo Ou and Hao Zhu for their assistance in C. elegans preparation, Li Yu and Dong Jiang for their assistance in Zebrafish embryo preparation, Wei Zhang and Yinjun Jia for their assistance in Drosophila larvae preparation, and Dan Zhang (Core Facility, Centre of Biomedical Analysis, Tsinghua University) for technical support with light-sheet microscopy. We would like to acknowledge the assistance of the Imaging Core Facility, Technology Centre for Protein Sciences, Tsinghua University, for assistance with Imaris 9.3 and Huygens. This work is supported by the National Natural Science Foundation of China (61327902, 61722209, 61729501, 61722209, and 61671265), the Beijing Municipal Science and Technology Commission (Z181100003118014), and the Natural Science Foundation of Beijing (18JQ019 and 7192103).

## Contributions

Q.D., and J.W. conceived and designed the project. J.W., J.F. and Z.L. designed and built the optical system. P.X. and J.W. designed the biological experiments. J.W., Z.L. and H.Q. conducted the numerical simulation and developed the reconstruction algorithm. T.Y. conducted the hardware synchronization. X.Z. and B.Z. performed cell cultures. Z.J. prepared the zebrafish embryos. Z.L., J.W., K.Z., X.L., T.Y., G.Z. and Z.J. conducted most biological experiments, data capturing and volume reconstruction. X.L. and L.F. conducted performance optimization on the algorithms. H.X. and J.F. conducted calibration and optimization of imaging setups. J.W., Z.L., J.F., P.X. and Q.D. prepared figures and wrote the manuscript with input from all authors.

## Materials & Correspondence

Correspondence and requests for materials should be addressed to (Q.D.), (P.X.), and (J.F.).

## References

1. Pantazis, P. & Supatto, W. Advances in whole-embryo imaging: A quantitative transition is underway. Nat. Rev. Mol. Cell Biol. 15, 327–339 (2014).

2. Liu, T.-L. et al. Observing the cell in its native state: Imaging subcellular dynamics in multicellular organisms. Science 360, eaaq1392 (2018).

3. Schulz, O. et al. Resolution doubling in fluorescence microscopy with confocal spinning-disk image scanning microscopy. Proc. Natl. Acad. Sci. U.S.A. 110, 21000–21005 (2013).

4. York, A. G. et al. Resolution doubling in live, multicellular organisms via multifocal structured illumination microscopy. Nat. Methods. 9, 749–754 (2012).

5. Schermelleh, L. et al. Subdiffraction Multicolor Imaging of the Nuclear Periphery with 3D Structured Illumination Microscopy. Science 320, 1332–1336 (2008).

6. Mertz, J. Optical sectioning microscopy with planar or structured illumination. Nat. Methods. 8, 811–819 (2011).

7. Ahrens, M. B., Orger, M. B., Robson, D. N., Li, J. M. & Keller, P. J. Whole-brain functional imaging at cellular resolution using light-sheet microscopy. Nat. Methods. 10, 413–420 (2013).

8. Bouchard, M. B. et al. Swept confocally-aligned planar excitation (SCAPE) microscopy for high-speed volumetric imaging of behaving organisms. Nat. Photonics. 9, 113–119 (2015).

9. Power, R. M. & Huisken, J. A guide to light-sheet fluorescence microscopy for multiscale imaging. Nat. Methods. 14, 360–373 (2017).

10. Winter, P. W. & Shroff, H. Faster fluorescence microscopy: advances in high speed biological imaging. Curr. Opin. Chen. Biol. 20, 46–53 (2014).

11. Levoy, M., Ng, R., Adams, A., Footer, M. & Horowitz, M. Light field microscopy. ACM Trans. Graph. 25, 924–934 (2006).

12. Prevedel, R. et al. Simultaneous whole-animal 3D imaging of neuronal activity using light-field microscopy. Nat. Methods. 11, 727–730 (2014).

13. Abrahamsson, S. et al. Fast multicolor 3D imaging using aberration-corrected multifocus microscopy. Nat. Methods. 10, 60–63 (2012).

14. Yang, W. et al. Simultaneous Multi-plane Imaging of Neural Circuits. Neuron 89, 269–284 (2016).

15. Nöbauer, T. et al. Video rate volumetric Ca2+ imaging across cortex using seeded iterative demixing (SID) microscopy. Nat. Methods. 14, 811–818 (2017).

16. Cong, L. et al. Rapid whole brain imaging of neural activity in freely behaving larval zebrafish (Danio rerio). Elife 6, 60 (2017).

17. Zhu, S., Lai, A., Eaton, K., Jin, P. & Gao, L. On the fundamental comparison between unfocused and focused light field cameras. Appl. Opt. 57, A1–11 (2018).

18. Martínez-Corral, M. & Javidi, B. Fundamentals of 3D imaging and displays: a tutorial on integral imaging, light-field, and plenoptic systems. Adv. Opt. Photonics. 10, 512–566 (2018).

19. Descloux, A. et al. Combined multi-plane phase retrieval and super-resolution optical fluctuation imaging for 4D cell microscopy. Nat. Photonics. 12, 165–172 (2018).

20. Ji, N., Milkie, D. E. & Betzig, E. Adaptive optics via pupil segmentation for high-resolution imaging in biological tissues. Nat. Methods. 7, 141–147 (2010).

21. Ji, N. Adaptive optical fluorescence microscopy. Nat. Methods. 14, 374–380 (2017).

22. Park, J.-H., Kong, L., Zhou, Y. & Cui, M. Large-field-of-view imaging by multi-pupil adaptive optics. Nat. Methods. 14, 581–583 (2017).

23. Adie, S. G., Graf, B. W., Ahmad, A., Carney, P. S. & Boppart, S. A. Computational adaptive optics for broadband optical interferometric tomography of biological tissue. Proc. Natl. Acad. Sci. U. S. A. 109, 7175–7180 (2012).

24. Shemonski, N. D. et al. Computational high-resolution optical imaging of the living human retina. Nat. Photonics. 9, 440–443 (2015).

25. Broxton, M. et al. Wave optics theory and 3-D deconvolution for the light field microscope. Opt. Express 21, 25418–25439 (2013).

26. Zheng, G., Horstmeyer, R. & Yang, C. Wide-field, high-resolution Fourier ptychographic microscopy. Nat. Photonics. 7, 739–745 (2013).

27. Waller, L., Situ, G. & Fleischer, J. W. Phase-space measurement and coherence synthesis of optical beams. Nat. Photonics. 6, 474–479 (2012).

28. Humphry, M. J., Kraus, B., Hurst, A. C., Maiden, A. M. & Rodenburg, J. M. Ptychographic electron microscopy using high-angle dark-field scattering for sub-nanometre resolution imaging. Nat. Commun. 3, 730 (2012).

29. Yang, H. et al. Electron ptychographic phase imaging of light elements in crystalline materials using Wigner distribution deconvolution. Ultramicroscopy 180, 173–179 (2017).

30. Alonso, M. A. Wigner functions in optics: Describing beams as ray bundles and pulses as particle ensembles. Adv. Opt. Photonics. 3, 272–365 (2011).

31. Cohen, N. et al. Enhancing the performance of the light field microscope using wavefront coding. Opt. Express 22, 24817–24839 (2014).

32. Liu, H.-Y., Zhong, J. & Waller, L. Multiplexed phase-space imaging for 3D fluorescence microscopy. Opt. Express 25, 14986–14995 (2017).

33. Ikoma, H., Broxton, M., Kudo, T. & Wetzstein, G. A convex 3D deconvolution algorithm for low photon count fluorescence imaging. Sci. Rep. 8, 11489 (2018).

34. Lim, Y. T., Park, J. H., Kwon, K. C. & Kim, N. Resolution-enhanced integral imaging microscopy that uses lens array shifting. Opt. Express 17, 19253–19263 (2009).

35. Llavador, A., Sánchez-Ortiga, E., Barreiro, J. C., Saavedra, G. & Martínez-Corral, M. Resolution enhancement in integral microscopy by physical interpolation. Biomed. Opt. Express 6, 2854–2863 (2015).

36. Nakamura, T., Horisaki, R. & Tanida, J. Computational phase modulation in light field imaging. Opt. Express 21, 29523–29543 (2013).

37. Boyd, S., Parikh, N., Chu, E., Peleato, B. & Eckstein, J. Distributed optimization and statistical learning via the alternating direction method of multipliers. Foundations and Trends in Machine Learning 3, 1–122 (2010).

38. Li, H. et al. Fast, volumetric live-cell imaging using high-resolution light-field microscopy. Biomed. Opt. Express 10, 29–49 (2019).

39. Sage, D. et al. DeconvolutionLab2: An open-source software for deconvolution microscopy. Methods 115, 28–41 (2017).

40. Ma, Y., Li, D., Smith, Z. J., Li, D. & Chu, K. Structured illumination microscopy with interleaved reconstruction (SIMILR). J. Biophotonics. 11, e201700090 (2017).

41. Huang, X. et al. Fast, long-term, super-resolution imaging with Hessian structured illumination microscopy. Nat. Biotechnol. 36, 451–459 (2018).

42. Chen, T.-W. et al. Ultrasensitive fluorescent proteins for imaging neuronal activity. Nature 499, 295–300 (2013).

43. Xu, J.-C. et al. Cultured networks of excitatory projection neurons and inhibitory interneurons for studying human cortical neurotoxicity. Sci. Transl. Med. 8, 333ra48 (2016).

44. Zhang, W., Yan, Z., Jan, L. Y. & Jan, Y. N. Sound response mediated by the TRP channels NOMPC, NANCHUNG, and INACTIVE in chordotonal organs of Drosophila larvae. Proc. Natl. Acad. Sci. U.S.A. 110, 13612–13617 (2013).

45. Shechtman, Y., Sahl, S. J., Backer, A. S. & Moerner, W. E. Optimal Point Spread Function Design for 3D Imaging. Phys. Rev. Lett. 113, 133902 (2014).

46. Thibault, P. & Menzel, A. Reconstructing state mixtures from diffraction measurements. Nature 494, 68–71 (2013).

47. Tokunaga, M., Imamoto, N. & Sakata-Sogawa, K. Highly inclined thin illumination enables clear single-molecule imaging in cells. Nat. Methods. 5, 159–161 (2008).

48. Ng, R. Fourier slice photography. ACM Trans. Graph. 24, 735–744 (2005).

49. Chai, Y. et al. Apoptotic regulators promote cytokinetic midbody degradation in C. Elegans. J. Cell Biol. 199, 1047–1055 (2012).

