## Supplementary Information for "3D observation of large-scale subcellular dynamics *in vivo* at the millisecond scale"

| | |
|---|---|
| **Extended Data Figure 1** | Concept and schematic design of scanning light field microscopy. |
| **Extended Data Figure 2** | The schematic diagram for the PSF modeling in phase space of sLFM. |
| **Extended Data Figure 3** | The comparisons of the PSFs and OTFs of WFM, direct aperture segmentation, and our DAOSLIMIT. |
| **Extended Data Figure 4** | The pipeline of the mutual iterative tomography algorithm with DAO. |
| **Extended Data Figure 5** | Comparison of the convergence speeds of traditional 3D deconvolution on simulated LFM data and our mutual iterative tomography on simulated sLFM data. |
| **Extended Data Figure 6** | Resolution characterization with different ratios of spatial overlap. |
| **Extended Data Figure 7** | Comparisons on fixed HeLa cells with GFP labeling on the actin and DAPI labeling on the nuclei. |
| **Extended Data Figure 8** | Performance comparison among DAOSLIMIT, commercial light-sheet microscopy, and commercial spinning-disk confocal microscopy. |
| **Extended Data Figure 9** | Motion artifacts elimination by the time-weighted algorithm on the imaging of histone-labeled C. elegans in vivo. |
| **Extended Data Figure 10** | Comparisons on the in vivo sample among WFM with 3D deconvolution, DAOSLIMIT without DAO, and DAOSLIMIT with DAO. |
| **Extended Data Figure 11** | The 3D calcium propagation along cultured rat cardiomyocytes at 100Hz. |
| **Supplementary Note 1** | Point spread function (PSF) of DAOSLIMIT. |
| **Supplementary Note 2** | Principle of incoherent synthetic aperture with the OTF analysis. |
| **Supplementary Note 3** | Principle of digital adaptive optics (DAO). |
| **Supplementary Note 4** | Mutual iterative tomography with DAO. |
| **Supplementary Table 1** | Imaging and reconstruction conditions for all fluorescence experiments. |
| **Supplementary Video 1** | 3D mitochondrial dynamics at 16Hz in a DRG neuron with DAO and without DAO. |
| **Supplementary Video 2** | 3D imaging of vesicles dynamics at 100Hz in one zebrafish epithelial cell at gastrulation stage. |
| **Supplementary Video 3** | Performance comparison deep inside a zebrafish embryos at gastrulation stage at 3Hz. |
| **Supplementary Video 4** | Vesicles and membrane morphological dynamics in zebrafish embryos at the gastrulation stage at 3Hz. |

| | |
|---|---|
| **Supplementary Video 5** | The dynamics of histone-labeled *C. elegans in vivo* at 16Hz. |
| **Supplementary Video 6** | Membrane dynamics in zebrafish embryos at the gastrulation stage *in vivo* at 10Hz. |
| **Supplementary Video 7** | Membrane dynamics in zebrafish embryos showing high-speed fluctuations of filopodia membrane. |
| **Supplementary Video 8** | The 3D calcium propagation in cultured rat cardiomyocytes at 100Hz. |
| **Supplementary Video 9** | Spontaneous 3D calcium propagation along a dendrite in a human 3D cerebral organoid infected with the GCamp6s indicators observed at 30Hz. |
| **Supplementary Video 10** | *In vivo* 3D calcium imaging of *Drosophila* larval Cho neurons at 100Hz with 500Hz sound stimulus. |

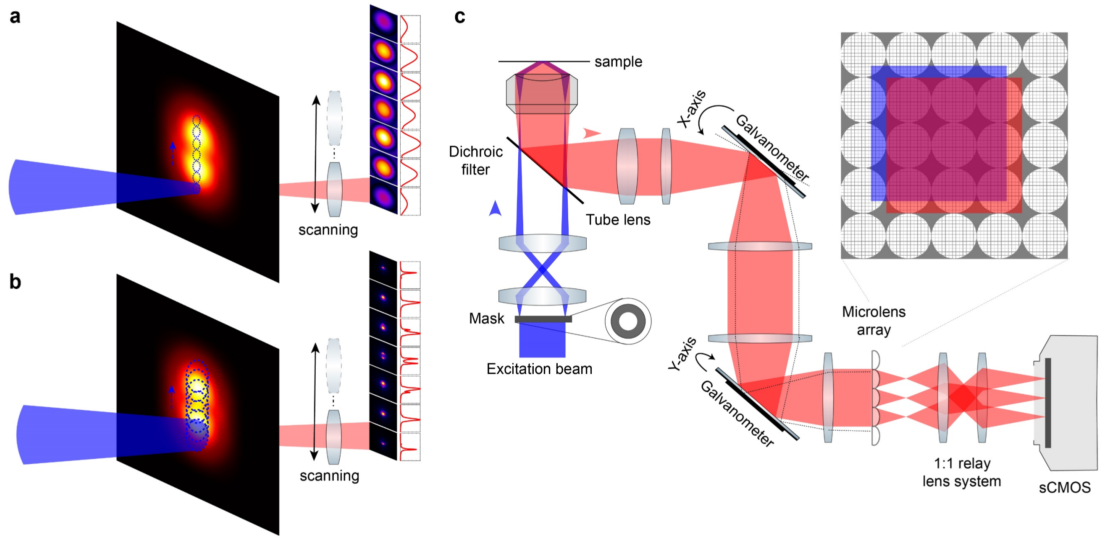


**Extended Data Figure 1 | Concept and schematic design of scanning light field microscopy. a**, Phase-space measurements of two point emitters at the defocus plane with an airy-unit aperture for spatial sampling, illustrating the low resolution in the spatial frequency domain due to the high-resolution spatial sampling. **b**, Overlapped phase-space measurements of the same sample with a larger aperture for spatial sampling to achieve high resolution in the spatial frequency domain. The spatial overlap helps to retrieve the compromised spatial resolution as the constraints for state unmixing, leading to high-resolution phase-space measurements. **c**, Schematic of the scanning light field microscopy system. A highly-inclined illumination excites the fluorescence within a small volume to reduce the out-of-focus background fluorescence and photo-induced damages. A microlens array is inserted on the image plane for parallel acquisition of low-resolution phase-space measurements. We use a two-dimensional galvo scanning system with a relay system between them to shift the image plane precisely and create the spatial overlap required for high-resolution multiplexed phase space, as illustrated in b. The inset on the upper-right corner illustrates the images on the microlens array before shifting(red) and after shifting(blue).

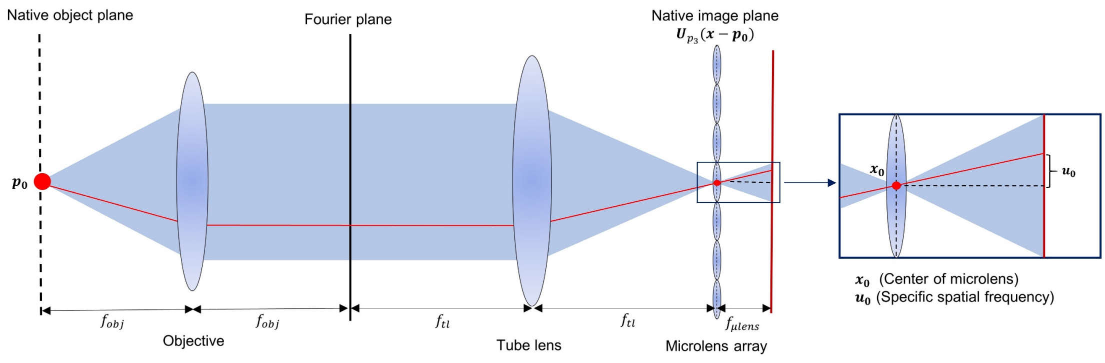


**Extended Data Figure 2 | The schematic diagram for the PSF modeling in phase space of sLFM.** An arbitrary 3D point with the lateral coordinates $\boldsymbol{p_0} = (p_1, p_2)$ and axial coordinate $p_3$ is placed at the sample plane. The complex field of the 3D point produces at the native image plane is $U_{p_3}(\boldsymbol{x} - \boldsymbol{p}_0)$. The center of the microlens is labeled as $\boldsymbol{x_0}$, while the pixels behind the microlens has a relative lateral displacement $\boldsymbol{u_0}$ to the center position.

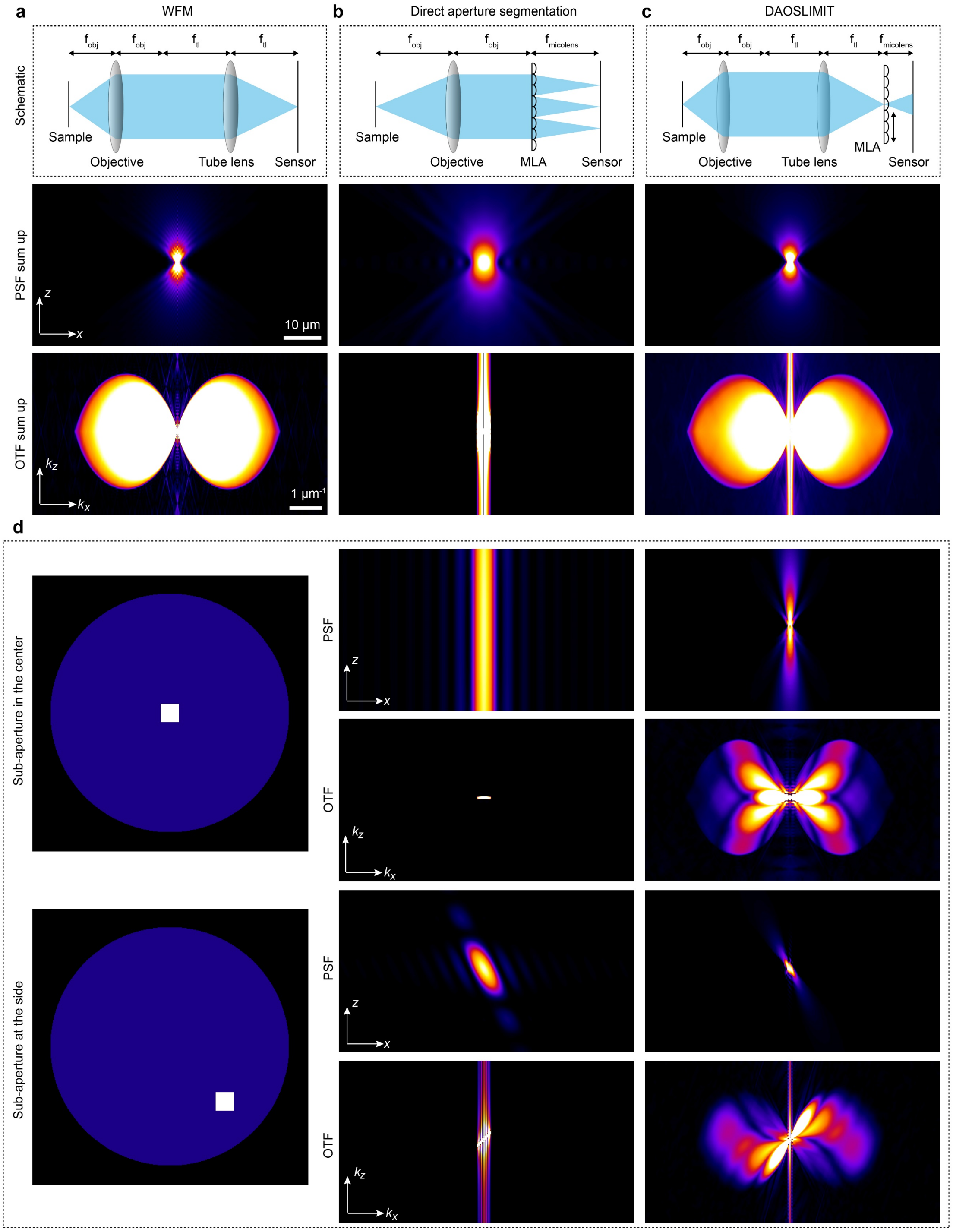


**Extended Data Figure 3 | The comparisons of the PSFs and OTFs of WFM, direct aperture segmentation, and our DAOSLIMIT. a-c**, The simulation is done based on the same objective 63×/1.4NA. Top row: illustrations of the different schematics. Second row: the sum-up of the PSFs of different frequency components. The PSF of WFM is used for comparison. Bottom row: the sum-up of the OTFs of different frequency components. The OTF of WFM is used for comparison. The sum-up OTF of our DAOSLIMIT can cover the same range as that of the WFM, illustrating the capability of the incoherent synthetic aperture with all the high-frequency information captured, while that of the scheme with direct aperture segmentation loses the high-frequency information. **d**, The PSFs and OTFs of two typical frequency components, whose corresponding sub-apertures are illustrated in the left panels.

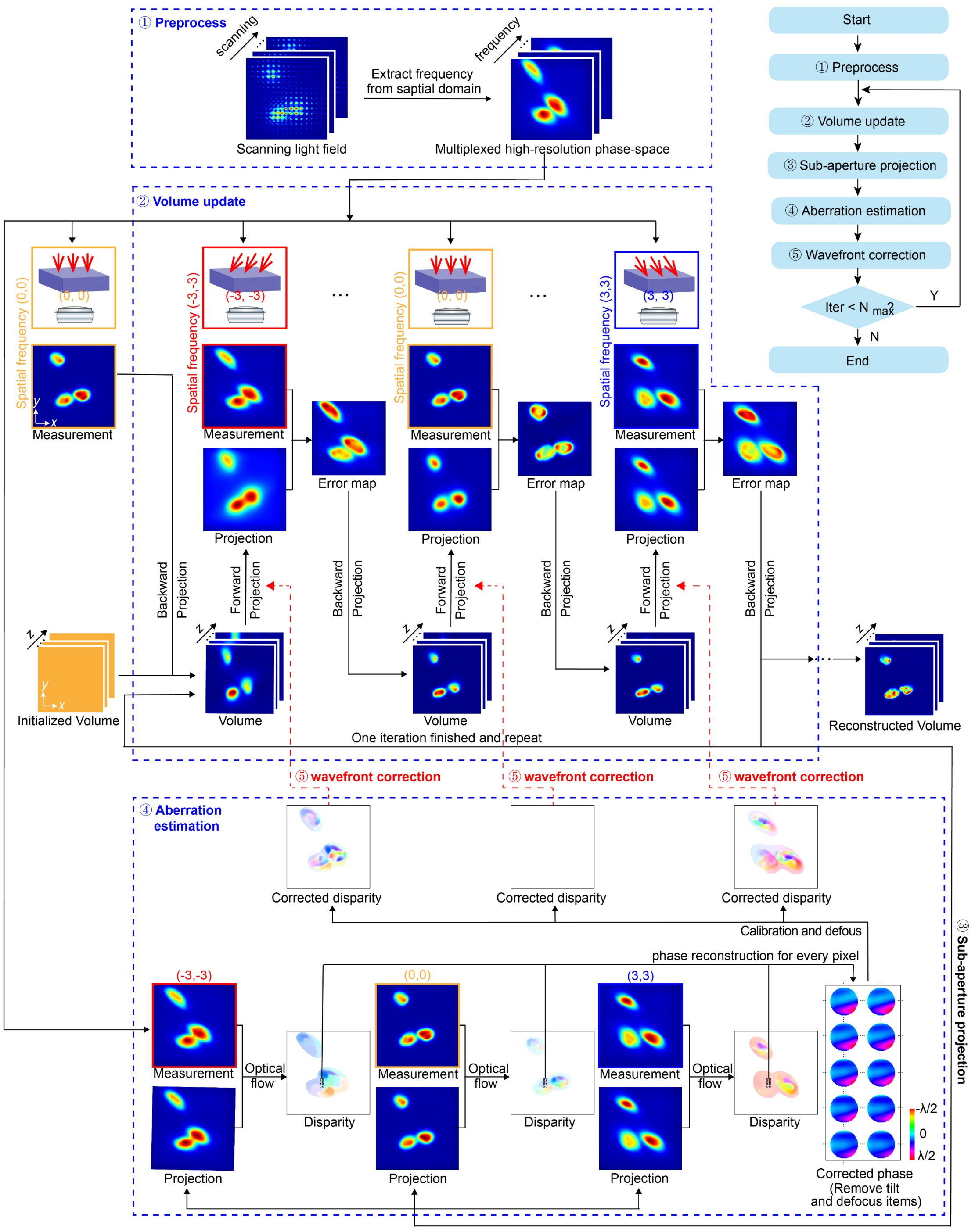


**Extended Data Figure 4 | The pipeline of the mutual iterative tomography algorithm with DAO.** First, high-resolution phase-space measurements were obtained with the pixel realignments of the sequentially scanned light field images. Secondly, a uniform volume is used as the initial volume for the start of iterations, which can be replaced by the results obtained from former frames in time-lapse imaging for faster convergence. To get the high-resolution volume with wavefront correction, the alternating direction method of multipliers is used to firstly reconstruct the 3D information with estimated aberration, and then estimate the optical aberration with the previous updated volume. During the volume update process, different spatial frequency components are used sequentially to update the high-resolution 3D volume, including both forward projections for error estimation and backward projections for correction. The correction wavefronts are incorporated in the forward projection step, which is set to zero for the first iteration. After the volume update, we employ a smooth optical flow algorithm to estimate the required disparity maps for the match between sub-aperture projections with non-corrected PSFs and the captured high-resolution phase-space data. After going through all the spatial frequency components, the disparity maps can be synthesized together to obtain a phase estimation for every pixel by simple integrals. For a continuous tiled reconstruction, we assume the aberration wavefront changes gently across the whole-FOV and remove the tilting and defocus components from the estimated wavefront of every pixel. The aberration wavefront for every pixel can then be corrected by applying different inverse shifts to different spatial frequency components during the forward projection in the next volume update.

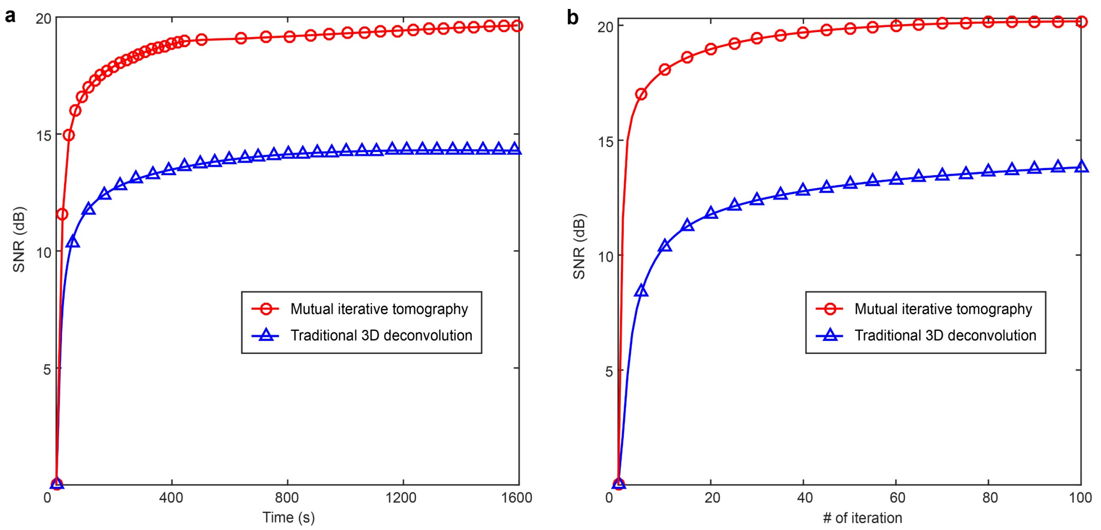


**Extended Data Figure 5 | Comparison of the convergence speeds of traditional 3D deconvolution on simulated LFM data and our mutual iterative tomography on simulated sLFM data. a**, The curves of the SNR versus computation time indicate that both algorithms require similar time to reach convergence, but our method has a much higher SNR due to the improved resolution. **b,** The curves of the SNR versus the number of iteration show the same result.

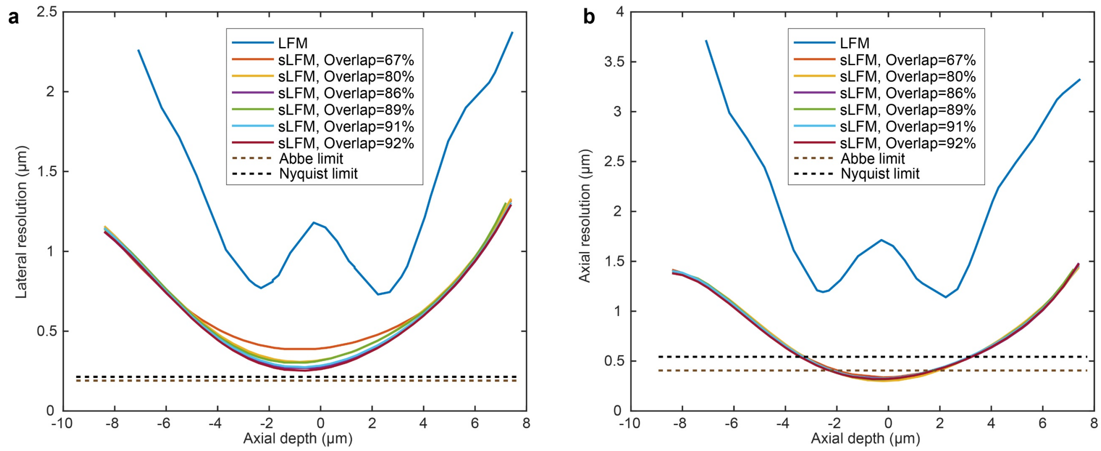


**Extended Data Figure 6 | Resolution characterization with different ratios of spatial overlap. a**, The lateral resolution of traditional LFM and sLFM with different spatial overlaps are characterized by sub-diffraction-limit fluorescence beads at different axial planes, indicating a similar performance with spatial overlap larger than 67%. **b**, The axial resolution of traditional LFM and sLFM with different spatial overlaps, indicating almost same performance with spatial overlap larger than 67%.

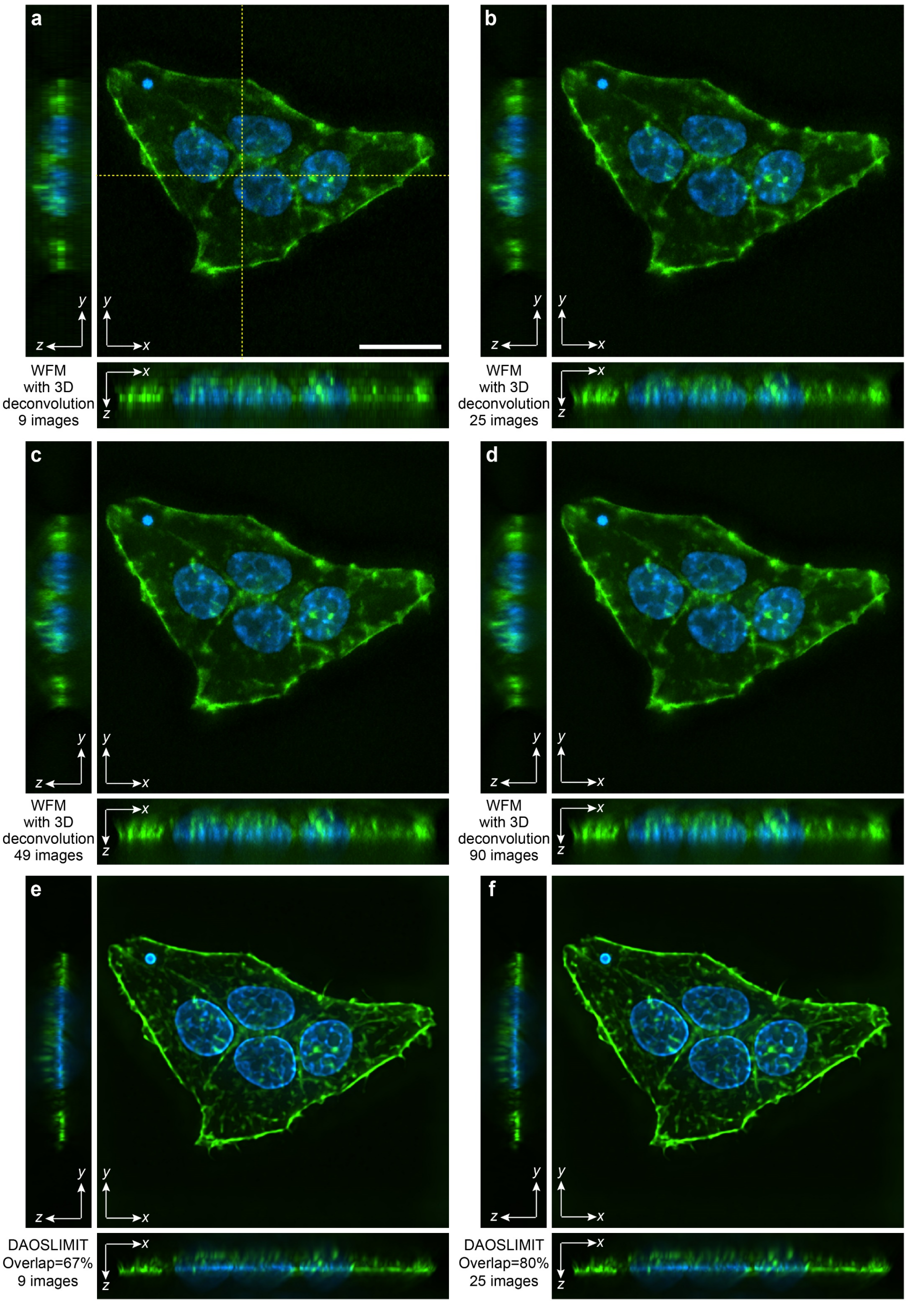


**Extended Data Figure 7 | Comparisons on fixed HeLa cells with GFP labeling on the actin and DAPI labeling on the nuclei. a-d**, The reconstructed orthogonal maximum intensity projections(MIPs) of WFM with different numbers of axial slices used in 3D RL deconvolution. **e, f**, Orthogonal MIPs by DAOSLIMIT with different spatial overlaps. To achieve a fair comparison, the same 63×/1.4NA objective was used with the same exposure time for each image. Scale bar: 20 μm.

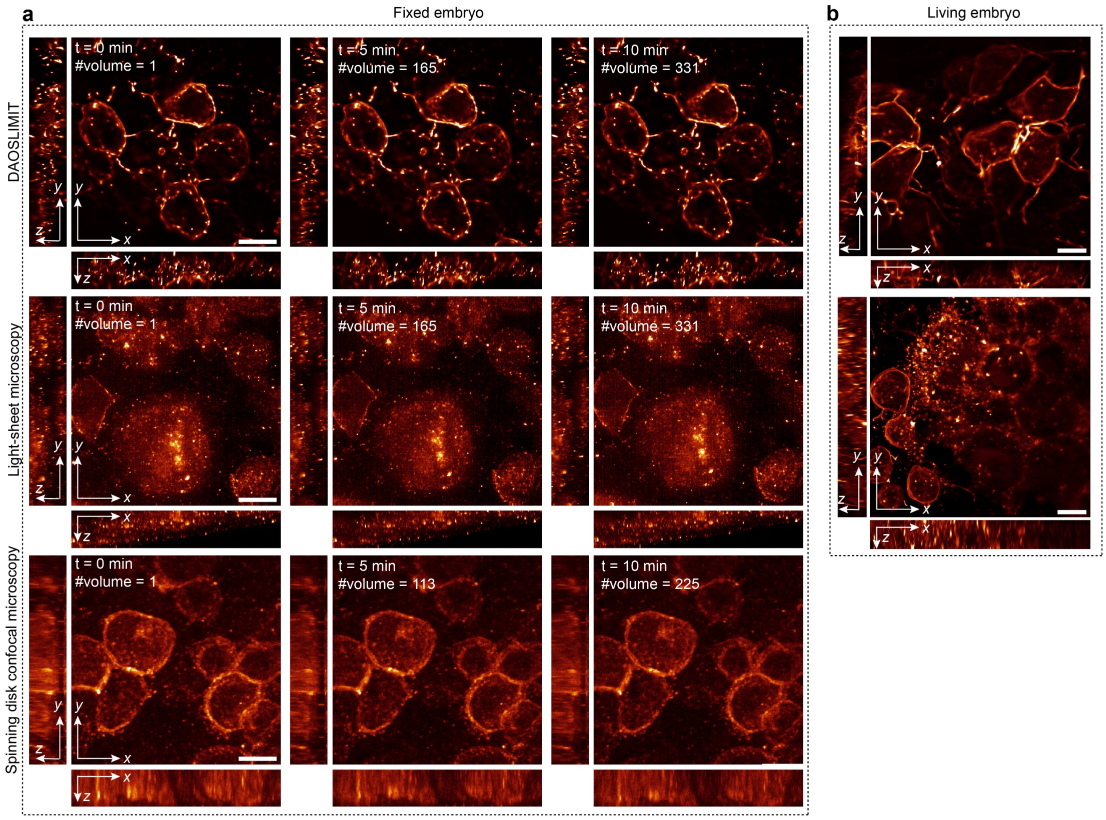


**Extended Data Figure 8 | Performance comparison among DAOSLIMIT, commercial light-sheet microscopy, and commercial spinning-disk confocal microscopy. a**, Time-lapse orthogonal MIPs of the fixed zebrafish gastrula obtained by DAOSLIMIT (Zeiss, 63×/1.4NA oil immersion objective), commercial light-sheet microscopy (Zeiss, Lightsheet Z.1, 63×/1.0NA water immersion objective) and spinning-disk confocal microscopy (PerkinElmer, Ultraview-Vox, Olympus, 60×/1.4NA oil immersion objective), respectively. The maximum 3D imaging speeds of the commercial microscopes were used with 50 axial layers and deconvolution were also applied to enhance the contrast. We kept imaging the sample for 10 minutes. The frame rate of our DAOSLIMIT was chosen to match the highest speed of the other two microscopes for a fair comparison. From the comparison, we can find that DAOSLIMIT can achieve the highest SNR with almost no photobleaching, while the other two microscopes both suffer from the scattering property of the fixed zebrafish gastrula, especially for the light-sheet microscope. The DAO capability provides such robustness to scattering. Scale bar: 10 μm. **b**, The orthogonal MIPs of the membrane-labeled living zebrafish embryo obtained by DAOSLIMIT and the light-sheet microscope, respectively. Due to the transparency of the living embryo, the performance of the light-sheet microscopy can be greatly enhanced, but the SNR has an apparent decrease from left to right (corresponding to the penetration of the excitation light sheet). However, DAOSLIMIT can achieve both high-resolution and high-SNR results with uniform performance. Scale bar: 10 μm.

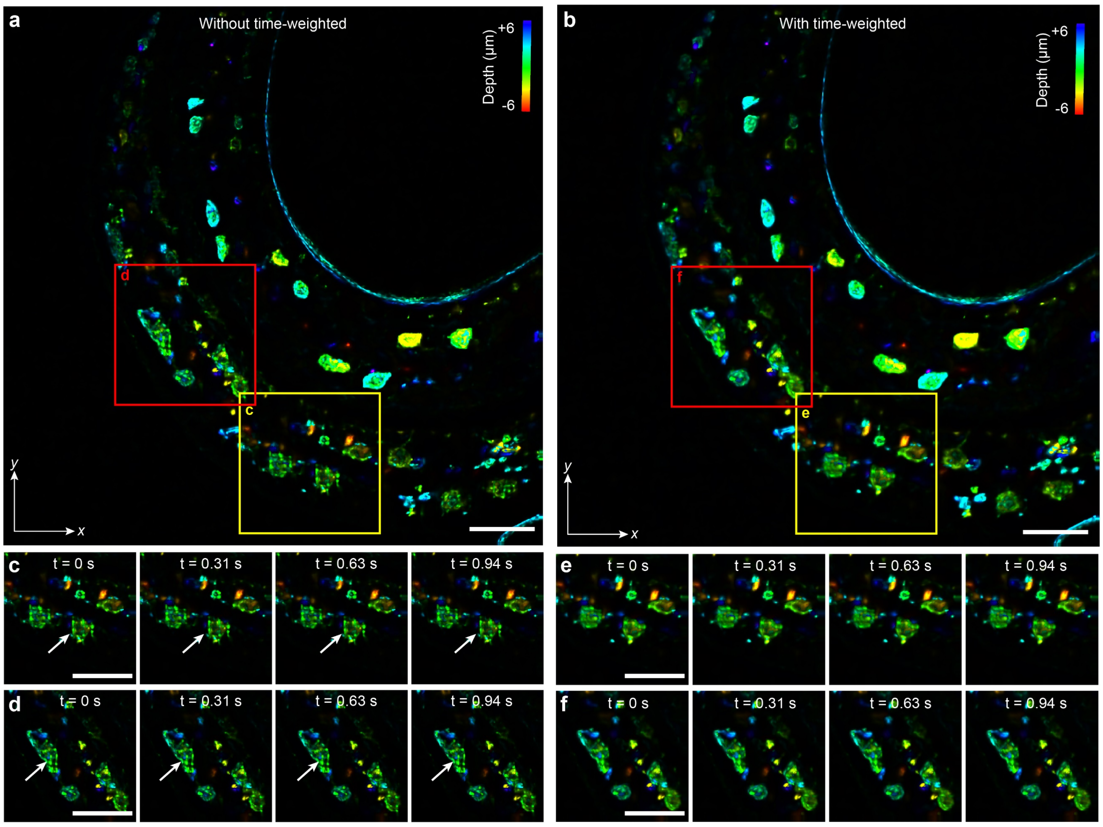


**Extended Data Figure 9 | Motion artifacts elimination by the time-weighted algorithm on the imaging of histone-labeled *C. elegans in vivo*.** **a**, Color-coded MIP of histone-labeled *C. elegans* without the time-weighted algorithm at t = 0 s. **b**, Color-coded MIP of histone-labeled *C. elegans* with the time-weighted algorithm at t = 0 s. **c**, Color-coded MIPs of the selected area (yellow box) in a at different time stamps, without the time-weighted algorithm. The white arrows point out the reconstruction artifacts due to high-speed motions. **d**, Color-coded MIPs of the selected area (red box) in a at different time stamps, without the time-weighted algorithm. The white arrows point out the reconstruction artifacts due to high-speed motions. **e**, Color-coded MIPs of the selected area (yellow box) in b at different time stamps, with the time-weighted algorithm. **f**, Color-coded MIPs of the selected area (red box) in b at different time stamps, with the time-weighted algorithm. All the motion artifacts are eliminated to show the effectiveness of the time-weighted algorithm, which can successfully compensate for the temporal resolution sacrificed by the scanning process in DAOSLIMIT. Scale bar: 10 μm.

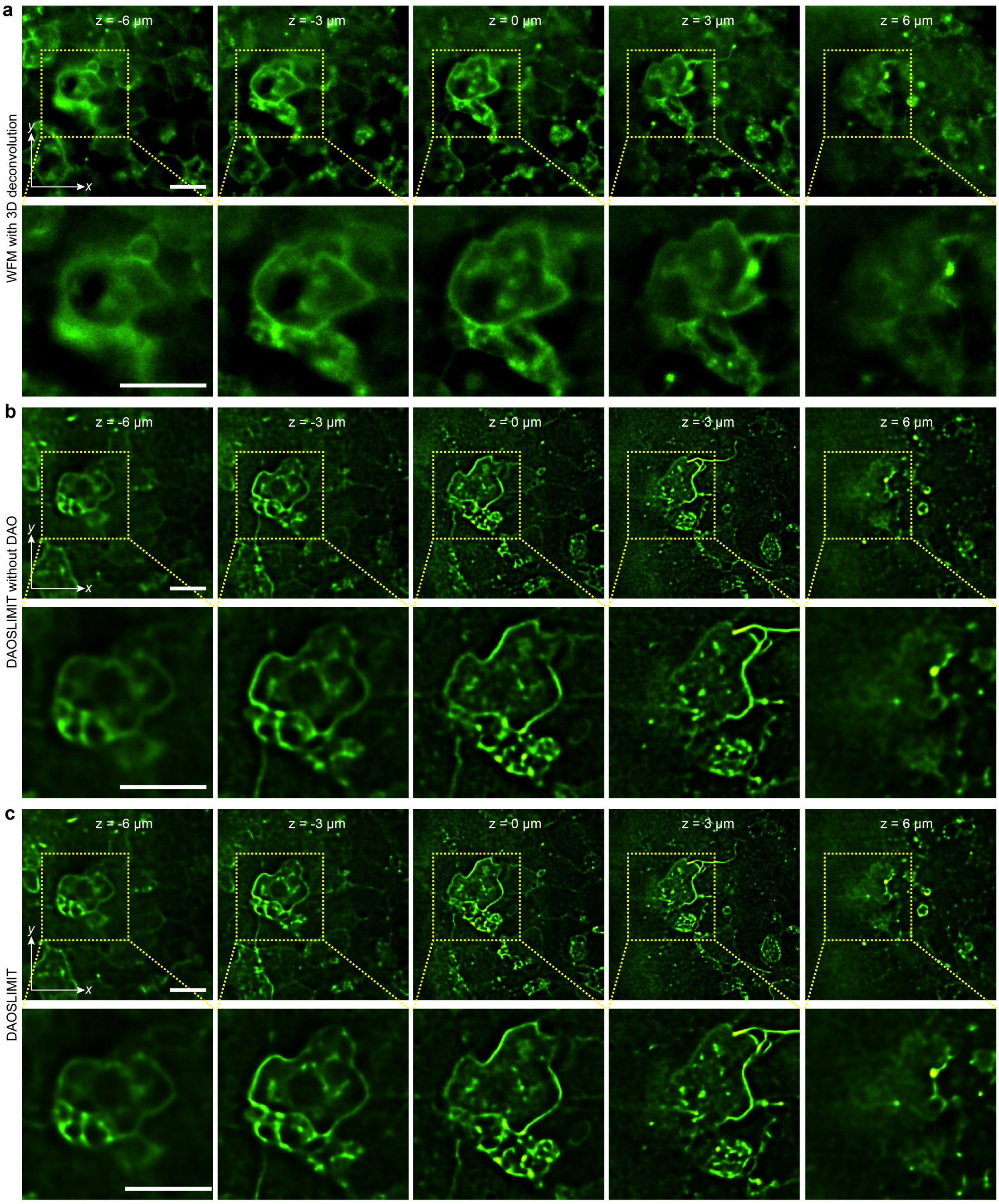


**Extended Data Figure 10 | Comparisons on the *in vivo* sample among WFM with 3D deconvolution, DAOSLIMIT without DAO, and DAOSLIMIT with DAO. a-c**, Different axial slices of the reconstructed membrane structures of the zebrafish embryo at gastrulation stage by WFM with 3D RL deconvolution, DAOSLIMIT without DAO and DAOSLIMIT with DAO, respectively. Zoom-in views with detailed comparisons show that substantial improvements in both SNR and resolution by DAOSLIMIT. Scale bar: 20 μm.

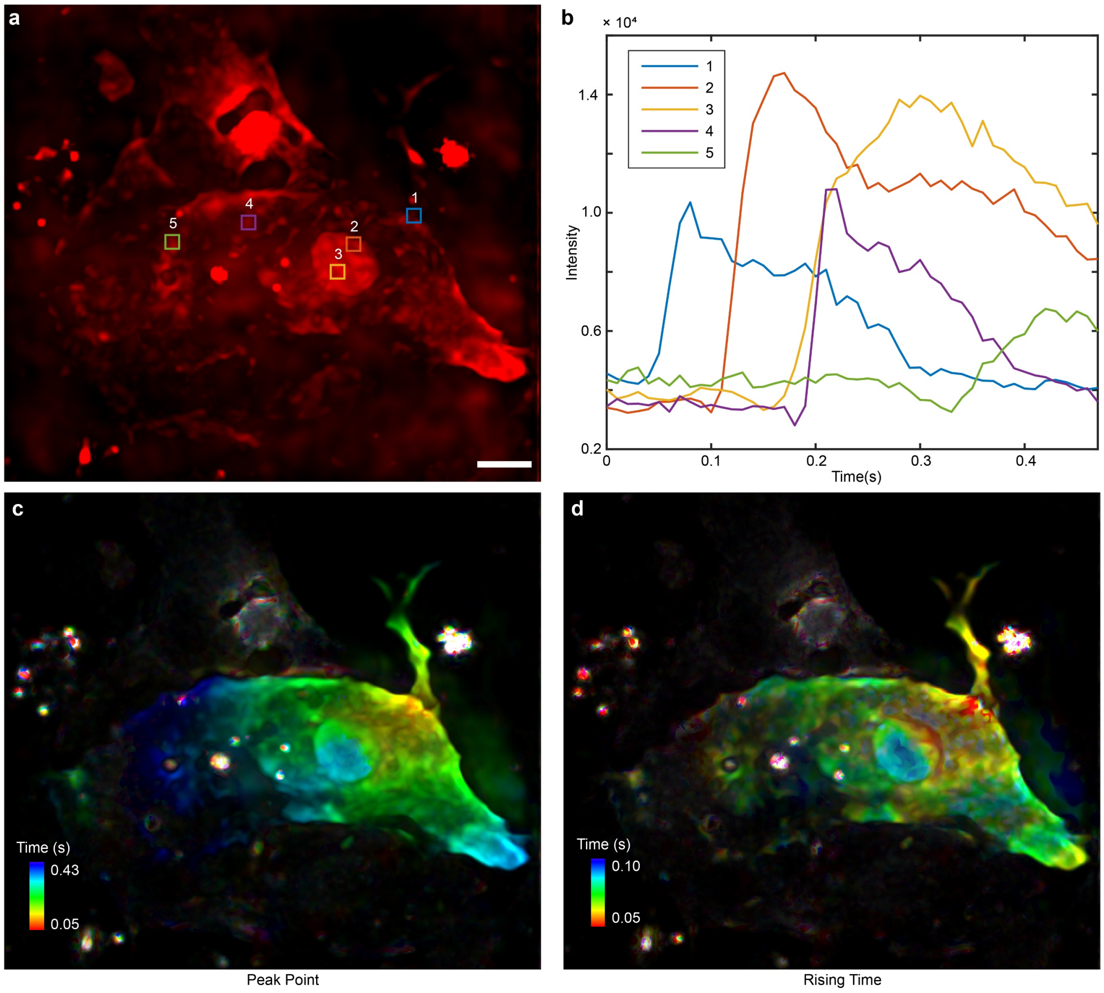


**Extended Data Figure 11 | The 3D calcium propagation along cultured rat cardiomyocytes at 100Hz. a**, The MIP of cardiac cells at t = 0.35 s. **b**, Different calcium response of five selected areas in a with curve fit for detailed analysis. **c**, The temporal-coded MIP of the calcium signals for peak time. Different colors visualize the peak instants of the calcium signal for every voxel. **d**, The temporal-coded MIP of the calcium signals for rising time. Different colors visualize the rising during (time required to increase from 20% to 80% of the maximum intensity) of the calcium signal. Scale bar: 10 µm.

### Supplementary Notes

#### Supplementary Note 1. Point spread function (PSF) of DAOSLIMIT

Light field microscopy (LFM) provides a way to measure the local variance of coherence with both angular information and spatial information within a snapshot. However, complicated spatially-nonuniform PSFs are used in traditional 3D deconvolution in LFM[25]. Reconstruction artifacts can be easily observed close to the native object plane. Such a problem is mainly owing to the low spatial resolution of the phase-space measurements by traditional LFM. Because of the fixed physical aperture of the microlens, traditional LFM can only sample the spatial domain at the step size equal to the pitch size of the microlens, which creates the spatially non-uniform PSFs with non-paraxial effects. The scanning process in DAOSLIMIT loosens the constraints and can achieve the high-resolution spatial sampling of the phase space with much smaller step sizes. Then we can model the imaging process of DAOSLIMIT in high-resolution multiplexed phase-space, which has a spatially-uniform and smooth PSF for different spatial frequency components.

All the denotations used here are labeled in Extended Data Fig. 2. Due to the incoherent property of fluorescence microscopy, we need to calculate the influence of an arbitrary 3D point with the lateral coordinates $\boldsymbol{p_0} = (p_1, p_2)$ and the axial coordinate $p_3$, to every sampling point by our DAOSLMIT (multiplexed phase-space measurements). Firstly, the complex field of the 3D point at the image plane can be formulated as below by Debye theory:

$$U_i(\boldsymbol{x}) = U_{p_3}(\boldsymbol{x}-\boldsymbol{p_0}) = \frac{M}{f_{obj}^2 \lambda^2} \exp\left(-i \cdot \frac{2\pi}{\lambda} p_3\right) \int_0^\alpha \sqrt{\cos\theta} \exp\left(-i \frac{4\pi p_3 \sin^2(\theta/2)}{\lambda}\right) J_0\left(\frac{2\pi \sin(\theta)}{\lambda} \sqrt{(x_1-p_1)^2+(x_2-p_2)^2}\right) \sin(\theta) d\theta \quad , \quad (1)$$

where $\boldsymbol{x} = (x_1, x_2)$ is the lateral coordinates on the image plane, while $M$ represents the magnification of the microscope, and $\lambda$ represents the wavelength of emission fluorescence. $f_{obj}$ is the focal length of the objective lens. $\alpha$ represents the halfangle of the numerical aperture. $J_0(\cdot)$ is the zeroth order Bessel function of the first kind.

Then we have a microlens array at the image plane, and the center position of the microlens is labeled as a vector $\boldsymbol{x_0}$. The modulation of the microlens with a center position $\boldsymbol{x_0}$ can be represented as below:

$$t(\boldsymbol{x}) = rect\left(\frac{\boldsymbol{x}-\boldsymbol{x_0}}{d_l}\right)\exp\left(\frac{-i\pi n}{\lambda f_{\mu lens}}\left\|\boldsymbol{x}-\boldsymbol{x_0}\right\|_2^2\right), \tag{2}$$

where $n$ is the refractive index of the sample, $f_{\mu lens}$ and $d_l$ represents the focal length and pitch size of a single microlens. $rect(\cdot)$ represents the 2D rectangle function, while $rect\left(\frac{\boldsymbol{x}-\boldsymbol{x_0}}{d_l}\right)$ illustrates the aperture of the microlens. Then the complex field $U_s$ at the back focal plane of this microlens, which is also the sensor plane, can be represented as below:

$$\begin{aligned}
U_s(\boldsymbol{x}') &= \frac{e^{j\frac{2\pi n}{\lambda}f_{\mu lens}}}{j\frac{2\pi n}{\lambda}f_{\mu lens}}\int_{\boldsymbol{x}} U_i(\boldsymbol{x})\cdot t(\boldsymbol{x})\cdot\exp\left\{j\frac{\pi n}{\lambda f_{\mu lens}}\left\|\boldsymbol{x}'-\boldsymbol{x}\right\|_2^2\right\}d\boldsymbol{x} \\
&= \frac{e^{j\frac{2\pi n}{\lambda}f_{\mu lens}}}{j\frac{2\pi n}{\lambda}f_{\mu lens}}\int_{\boldsymbol{x}} U_{p_3}(\boldsymbol{x}-\boldsymbol{p_0})\cdot rect\left(\frac{\boldsymbol{x}-\boldsymbol{x_0}}{d_l}\right)\exp\left(j\frac{-\pi n}{\lambda f_{\mu lens}}\left\|\boldsymbol{x}-\boldsymbol{x_0}\right\|_2^2\right)\cdot\exp\left\{j\frac{\pi n}{\lambda f_{\mu lens}}\left\|\boldsymbol{x}'-\boldsymbol{x}\right\|_2^2\right\}d\boldsymbol{x} \\
&= \frac{e^{j\frac{2\pi n}{\lambda}f_{\mu lens}}}{j\frac{2\pi n}{\lambda}f_{\mu lens}}\int_{\boldsymbol{x}} U_{p_3}(\boldsymbol{x}+\boldsymbol{x_0}-\boldsymbol{p_0})\cdot rect\left(\frac{\boldsymbol{x}}{d_l}\right)\exp\left(j\frac{-\pi n}{\lambda f_{\mu lens}}\left\|\boldsymbol{x}\right\|_2^2\right)\cdot\exp\left\{j\frac{\pi n}{\lambda f_{\mu lens}}\left\|(\boldsymbol{x}'-\boldsymbol{x_0})-\boldsymbol{x}\right\|_2^2\right\}d\boldsymbol{x} \\
&= \frac{e^{j\frac{2\pi n}{\lambda}f_{\mu lens}}}{j\frac{2\pi n}{\lambda}f_{\mu lens}}\exp\left(j\frac{\pi n}{\lambda f_{\mu lens}}\left\|\boldsymbol{x}'-\boldsymbol{x_0}\right\|_2^2\right)\int_{\boldsymbol{x}} U_{p_3}(\boldsymbol{x}+\boldsymbol{x_0}-\boldsymbol{p_0})\cdot rect\left(\frac{\boldsymbol{x}}{d_l}\right)\cdot\exp\left\{j\frac{2\pi n}{\lambda f_{\mu lens}}(\boldsymbol{x}'-\boldsymbol{x_0})^T\boldsymbol{x}\right\}d\boldsymbol{x} \\
&= \frac{e^{j\frac{2\pi n}{\lambda}f_{\mu lens}}}{j\frac{2\pi n}{\lambda}f_{\mu lens}}\exp\left(j\frac{\pi n}{\lambda f_{\mu lens}}\left\|\boldsymbol{x}'-\boldsymbol{x_0}\right\|_2^2\right)F_{\boldsymbol{\omega}}\left(U_{p_3}(\boldsymbol{x}+\boldsymbol{x_0}-\boldsymbol{p_0})\cdot rect\left(\frac{\boldsymbol{x}}{d_l}\right)\right)
\end{aligned}, \tag{3}$$

where $\boldsymbol{x}'$ represents the lateral coordinates at the sensor plane after the Fresnel propagation, $F_{\omega}(\cdot)$ is the 2D Fourier transform function, where $\boldsymbol{\omega} = \frac{2\pi n}{\lambda f_{\mu lens}}(\boldsymbol{x}'-\boldsymbol{x_0})$.

Different pixels behind the microlens with a center position $\boldsymbol{x_0}$ has a relative lateral displacement $\boldsymbol{u_0}$ to the center position, as shown in Extended Data Fig. 2, which also corresponds to different spatial frequency. The sampling of the sensor pixel can then be defined as another rectangle function with an aperture size of the sensor pixel size $d_s$:

$$s(\boldsymbol{x}) = rect\left(\frac{\boldsymbol{x}' - \boldsymbol{x_0} - \boldsymbol{u_0}}{d_s}\right) \quad . \tag{4}$$

Above all, the influence of an arbitrary 3D point with the axial coordinate $p_3$ to every measurement in multiplexed phase-space can be represented as below:

$$\begin{aligned} W_{p_3}(\boldsymbol{x_0}, \boldsymbol{p_0}, \boldsymbol{u_0}) &= \int_{\boldsymbol{x}'} \left\| U_s(\boldsymbol{x}') \cdot s(\boldsymbol{x}) \right\|_2^2 d\boldsymbol{x}' \\ &= \int_{\boldsymbol{x}'} \left\| \frac{e^{j\frac{2\pi n}{\lambda} f_{\mu lens}}}{j\frac{2\pi n}{\lambda} f_{\mu lens}} \exp\left( j\frac{\pi n}{\lambda f_{\mu lens}} \left\| \boldsymbol{x}' - \boldsymbol{x_0} \right\|_2^2 \right) F_{\omega}\left( U_{p_3}(\boldsymbol{x} + \boldsymbol{x_0} - \boldsymbol{p_0}) \cdot rect\left(\frac{\boldsymbol{x}}{d_l}\right) \right) \cdot rect\left( \frac{\boldsymbol{x}' - \boldsymbol{x_0} - \boldsymbol{u_0}}{d_s} \right) \right\|_2^2 d\boldsymbol{x}' \\ &\xlongequal{\boldsymbol{x}'' = \boldsymbol{x}' - \boldsymbol{x_0}} \int_{\boldsymbol{x}''} \left\| \frac{e^{j\frac{2\pi n}{\lambda} f_{\mu lens}}}{j\frac{2\pi n}{\lambda} f_{\mu lens}} \exp\left( j\frac{\pi n}{\lambda f_{\mu lens}} \left\| \boldsymbol{x}'' \right\|_2^2 \right) F_{\frac{2\pi n}{\lambda f_{\mu lens}}\boldsymbol{x}''}\left( U_{p_3}(\boldsymbol{x} + \boldsymbol{x_0} - \boldsymbol{p_0}) \cdot rect\left(\frac{\boldsymbol{x}}{d_l}\right) \right) \cdot rect\left( \frac{\boldsymbol{x}'' - \boldsymbol{u_0}}{d_s} \right) \right\|_2^2 d\boldsymbol{x}'' \end{aligned} \tag{5}$$

From the representation of the multiplexed phase space measurement, we can clearly observe the 2D spatial-invariance property for different spatial frequency components:

$$W_{p_3}(\boldsymbol{x_0} + \Delta\boldsymbol{x}, \boldsymbol{p_0} + \Delta\boldsymbol{x}, \boldsymbol{u_0}) = W_{p_3}(\boldsymbol{x_0}, \boldsymbol{p_0}, \boldsymbol{u_0}) \ , \tag{6}$$

Such a property can not only accelerate the reconstruction algorithm but also divide the imaging process into different segmented spatial frequency components, which provides the basis for digital adaptive optics. In this case, the PSF $H(\boldsymbol{x_0}, p_3, \boldsymbol{u_0})$ of the imaging process modeled in phase space can be represented as:

$$H(\boldsymbol{x_0}, p_3, \boldsymbol{u_0}) = W_{p_3}(\boldsymbol{x_0}, \boldsymbol{0}, \boldsymbol{u_0}), \tag{7}$$

Then the imaging process of 3D fluorescence sample $V(\boldsymbol{x}, z)$ with a 2D lateral coordinate $\boldsymbol{x}$ and an axial coordinate $z$, can be illustrated as below:

$$M(\boldsymbol{x}, \boldsymbol{u}) = \int_z V(\boldsymbol{x}, z) * H(\boldsymbol{x}, z, \boldsymbol{u}) dz, \tag{8}$$

where $\boldsymbol{u}$ represents the 2D coodinates in the spatial frequency domain, and $M\left(\boldsymbol{x},\boldsymbol{u}\right)$ is the multiplexed phase-space measurements captured by the scanning LFM.

**Supplementary Note 2. Principle of incoherent synthetic aperture with the OTF analysis**

Synthetic aperture is a powerful technique applied in various coherent imaging modes such as radar and optical coherence tomography. However, it requires coherence between small apertures and is hard to be employed in incoherent imaging modes such as fluorescence microscopy by simply segmenting the aperture plane due to the loss of phase information. The principle of the incoherent synthetic aperture by our DAOSLIMIT is the addition of coherence by frequency aliasing.

The small aperture of each microlens $d_l$ (100 μm in our system), which is comparable to the diffraction-limit at the image plane (~13.7 μm at 500nm), creates frequency aliasing between different spatial frequency components. As indicated by Eq. 5, the spatial frequency components obtained from the local spatial sampling are different from those obtained from the direct aperture segmentation, because the dot product and the Fourier transform in Eq. 5 cannot change the order.

To show the difference in detail, we simulated the OTFs of different schemes, including wide-field fluorescence microscopy (WFM), direct aperture segmentation (with the lens array placed at the pupil plane) and our DAOSLIMIT (with lens array placed at the image plane and the lateral scanning capability for dense sampling). The results are shown in Extended Data Fig. 3. The PSF and OTF of WFM are used for comparison, showing the diffraction-limited performance without 3D imaging capability. Then we sum up the PSFs and OTFs of different spatial-frequency components of the scheme with direct aperture segmentation and our DAOSLIMIT for comparison. All the schemes have the same light efficiency without light loss, meaning that we can scale the sum-up PSF based on the same total intensity for a fair comparison. The OTFs are shown on the logarithmic scale. With the strong frequency aliasing introduced by the small microlens aperture, the sum-up OTF of our DAOSLIMIT covers the same frequency range as the WFM's OTF. Therefore, DAOSLIMIT can achieve diffraction-limited incoherent synthetic aperture. But DAOSLIMIT has a much larger depth-of-field for 3D imaging capability. However, the sum-up OTF of the scheme with direct aperture segmentation covers a much smaller region, indicating the loss of high-frequency information.

The PSFs and OTFs of several spatial frequency components are also shown in the Extended Data Fig. 3d, illustrating that the frequency aliasing in DAOSLIMIT extends the OTFs of different spatial frequency component, which works as the coherence between them and is essential for synthetic aperture. The OTFs of different spatial frequency components emphasize different areas in the spatial frequency domain, which makes them more robust to noise. In the meantime, the spatial frequency components of the scheme with direct aperture segmentation lose the coherence and have only low-frequency information left.

**Supplementary Note 3. Principle of digital adaptive optics (DAO)**

Optical aberrations induced by the optical system and imaging environment will reduce the resolution and SNR in 3D fluorescence imaging, especially for high-NA objectives. Various adaptive optics techniques[21] are proposed to correct the wavefront aberration and improve imaging performance in scattering tissues and multicellular organisms. A common way[2,20] is to segment the rear pupil function into subregions, and deflect each group of rays toward focus by a deformable mirror[20] or spatial light modulator[22] with a good estimation of the aberrated wavefront. However, such systems need either a wavefront sensor or iterative searching process with a guidestar, which compromises the imaging speed and system simplicity.

Our DAOSLIMIT provides a new computational way to estimate and correct the aberration in post-processing with the high-resolution multiplexed phase-space measurements. According to the translation-shifting property of Fourier theorem, the lateral shift distance $\boldsymbol{a}$ of a complexed field is equivalent to applying a linear phase modulation $\varphi(\boldsymbol{\omega_x}) = \exp\left(-i2\pi\boldsymbol{\omega_x a}\right)$ at the back-pupil plane, which can be represented as below:

$$F_\omega\left(H(\boldsymbol{x}-\boldsymbol{a})\right) = F_\omega\left(H(\boldsymbol{x})\right)\cdot\exp\left(-i2\pi\boldsymbol{\omega_x a}\right), \tag{9}$$

where $H(\boldsymbol{x})$ represents an arbitrary complex field.

Similar to the traditional adaptive optics with the deformable mirror array, the correction phase can be approximated by linear fitting of segmented pupils. Then, the spatial shift of different spatial frequency components $\boldsymbol{u_0}$ can be approximately regarded as a phase gradient at the corresponding sub-apertures. In this case, we can simply shift the sub-aperture PSFs during reconstruction to apply an approximate wavefront correction to the system.

Due to the optical heterogeneity of multicellular organisms, the aberrations at different areas across a large FOV should be estimated separately. In practice, we calculate the lateral shifts of each sub-aperture PSF by a smooth optical flow algorithm, with an assumption that the aberrations change smoothly within the whole FOV.

**Supplementary Note 4. Mutual iterative tomography with DAO**

The pipeline of the algorithm is shown in the Extended Data Fig. 4. Firstly, high-resolution phase-space measurements were obtained with the pixel realignments of the sequentially-scanned light field images, according to Eq. 5. The sensor pixels after each microlens with the same relative distance $\boldsymbol{u}$ to the center of the microlens, are selected out as a specific spatial frequency $\boldsymbol{u}$. These pixels are arranged together according to the spatial positions of their corresponding microlens, whose center position is $\boldsymbol{x}$. Without scanning just like traditional LFM, the minimum sampling interval of $\boldsymbol{x}$ is limited by the pitch size of the microlens $d_l$. However, our scanning process in DAOSLIMIT can increase the sampling rate. Therefore, all the pixels of the sequentially captured light field images are realigned in the multiplexed 4D phase space according to their $\boldsymbol{x}$ and $\boldsymbol{u}$ to get the multiplexed phase-space measurements $M(\boldsymbol{x},\boldsymbol{u})$.

When the overlap ratio is smaller than 92% (corresponding to 13×13 lateral shifts), we will conduct a cubic interpolation to upsample the $M(\boldsymbol{x},\boldsymbol{u})$ to keep the voxel size small enough for the resolution improvement in reconstruction due to aperture synthesis. The experiments on fluorescence beads and biological samples (Extended Data Fig. 8 and 9) illustrate that the overlap ratio of 67%, corresponding to 3×3 lateral shifts, is sufficient for diffraction-limited performance in most conditions.

Secondly, a uniform volume $\boldsymbol{g^0}$ is used as the initial volume for the start of iterations, which can be replaced by the results obtained from former frames in time-lapse imaging for faster convergence. We use the ADMM algorithm[37] to update the volume and aberration iteratively. During every volume update, different spatial frequency components (from $\boldsymbol{u}_1$ to $\boldsymbol{u}_N$) are used sequentially to achieve the high-resolution 3D volume, including both forward projections for error estimation and backward projections for correction. For each spatial frequency components $\boldsymbol{u}_j$, the forward projection can be formulated as below:

$$P(\boldsymbol{x}) = \int_z g_{j-1}^k(\boldsymbol{x}, z) * H(\boldsymbol{x}, z, \boldsymbol{u}_j) dz, \tag{10}$$

where $g_{j-1}^k(\boldsymbol{x}, z)$ is the volume updated from the last frequency components (specifically we define $g_0^k(\boldsymbol{x}, z) = g_N^{k\text{-}1}(\boldsymbol{x}, z)$), and $*$ represents the 2D convolution process in the lateral domain (only applied to $\boldsymbol{x}$). Then we shift the forward projection with the disparity map $disp^{k\text{-}1}(\boldsymbol{x}, \boldsymbol{u}_j)$ according to the correction wavefront estimated from the last iteration (which is set to zeros for the first iteration):

$$P_{corr}(\boldsymbol{x}) = P\left(\boldsymbol{x} + disp^{k\text{-}1}(\boldsymbol{x}, \boldsymbol{u}_j)\right). \tag{11}$$

Such a process can be regarded as the wavefront correction digitally. The error can be calculated by the comparison with the measurements as below:

$$Error(\boldsymbol{x}) = M(\boldsymbol{x}, \boldsymbol{u}_j) / P_{corr}(\boldsymbol{x}), \tag{12}$$

which can be back-propagated to get the gradient for volume update:

$$G_{corr}(\boldsymbol{x}, z) = Error(\boldsymbol{x}) * H^T(\boldsymbol{x}, z, \boldsymbol{u}_j). \tag{13}$$

The new volume estimated in this iteration can be represented as:

$$g_{j+1}^k(\boldsymbol{x}, z) \leftarrow w_{\boldsymbol{u}_j} g_j^k(\boldsymbol{x}, z) \odot G_{corr}(\boldsymbol{x}, z) / \left(J * H^T(\boldsymbol{x}, z, \boldsymbol{u}_j)\right) + \left(1 - w_{\boldsymbol{u}_j}\right) g_j^k(\boldsymbol{x}, z), \tag{14}$$

where $\odot$ represents the dot product process, $J$ represents the all-ones matrix, and $w_{\boldsymbol{u}_j}$ is the weight used to balance the different shot noise for different spatial frequency components. The $w_{\boldsymbol{u}_j}$ is calculated based on the energy distribution of the PSF:

$$w_{\boldsymbol{u}_j} = c \frac{\left\| H(\boldsymbol{x}, z, \boldsymbol{u}_j) \right\|_1}{\sum_{\boldsymbol{u}_k = \boldsymbol{u}_1}^{\boldsymbol{u}_N} \left\| H(\boldsymbol{x}, z, \boldsymbol{u}_k) \right\|_1}, \tag{15}$$

where $c$ is the coefficient to balance the convergence rate and performance, which depends on the number of frequency components. We choose $c = 80$ for all of our experiments because the number of frequency components in our setup is 169.

After going through all the spatial frequency components with aberration correction, we fix the updated volume and use it to estimate the aberration wavefronts.

We employ a smooth optical flow algorithm to estimate the required disparity maps for the match between sub-aperture projections with the captured high-resolution phase-space measurements. For each spatial frequency component $\boldsymbol{u}_j$, we can get the disparity map by a smooth optical flow estimation between the forward projection $P_j(\boldsymbol{x}) = \int_z g_{j-1}^k(\boldsymbol{x}, z) * H(\boldsymbol{x}, z, \boldsymbol{u}_j) dz$ and the phase space measurements $M(\boldsymbol{x}, \boldsymbol{u}_j)$:

$$disp_{temp}^k(\boldsymbol{x}, \boldsymbol{u}_j) = OF(P_j(\boldsymbol{x}), M(\boldsymbol{x}, \boldsymbol{u}_j)). \tag{16}$$

where $OF(I_a, I_b)$ is a function to calculate the optical flow between $I_a$ and $I_b$. In our prototype, we calculate the optical flow by finding the maximum correlation positions for several segmented regions (usually 7×7). For a continuous tiled reconstruction, we assume the aberration wavefront changes gently across the whole-FOV. Therefore, we use a cubic interpolation to create the initial disparity map for every pixel based on the disparities estimated from segmented regions. After going through all the frequency components, the disparity maps can be synthesized together to obtain different phase estimations for every pixel. We further remove the tilting and defocus components from the estimated wavefront of every pixel, and obtain the new disparity map $disp^k(\boldsymbol{x}, \boldsymbol{u}_j)$ for next volume update with aberration correction.

The whole algorithm usually takes about 5 to 10 iterations for convergence, as shown in Extended Data Fig. 7. And only 2 iterations are required for time-lapse video with our time-loop algorithms illustrated before. The actual time required for the volume reconstruction has quite a large variation due to the volume size, iteration numbers and the working stations used. For a typical, one iteration of a volume with around 800×800×119 voxels takes about 50s with one NVIDIA GeForce GTX 1080Ti by our proof-of-concept implementations based on Matlab 2018.

**Supplementary Table 1. Imaging and reconstruction conditions for all fluorescence experiments.**

| | **Sample, (imaging T, ℃)** | **Fluorescent label** | **Cropped voxel volume (dx×dy ×dz)** | **Cropped image Volume (x,y,z) $\mu m^3$** | **Exposure time (# time pts)** | **λ: Power ($mW/cm^2$)** | **Volume rate (Hz)** |
|---|---|---|---|---|---|---|---|
| 2a, ED6 | Fluorescence beads 37℃ | Yellow-green fluorescent (505/515) | 273×273×97 | 30.0×30.0 ×19.4 | 50ms 33pts | 488: (28) | - |
| 2c-f | Fluorescence beads 37℃ | Yellow-green fluorescent (505/515) | 793×793×97 | 87.2×87.2 ×19.4 | 50ms 1pts | 488: (28) | - |
| 3, ED7 | Hela cell 37℃ | DAPI GFP | 923×923×81 | 101.5×101.5×16.2 | 50ms 1pts | 454: (4.2) 488: (35) | - |
| 4, SV1 | Mitochondria-labelled neuron 37℃ | EGFP | 819×819×81 | 90.1×90.1 ×16.2 | 50ms 3823pts | 488: (28) | 16 |
| 5a | zebrafish embryo membrane | EGFP | 1053×1053×119 | 115.8×115.8×23.8 | 50ms 1pts | 488: (16) | - |
| 5b, SV2 | zebrafish embryo vesicles | EGFP | 611×611×119 | 67.2×67.2 ×23.8 | 5ms 2000pts | 488: (35) | 100 |
| 5c-g, SV3, SV4 | zebrafish embryo membrane | EGFP | 1053×1053×119 | 115.8×115.8×23.8 | 300ms 4800pts | 488: (35) | 3 |
| 5h, ED10, SV6 | zebrafish embryo membrane | EGFP | 1001×1001×119 | 110.1×110.1×23.8 | 80ms 5000pts | 488: (35) | 10 |

| | | | | | | | |
|---|---|---|---|---|---|---|---|
| 5i, SV7, ED8b | zebrafish embryo membrane | EGFP | 1079×1079×119 | 118.7×118.7×23.8 | 80ms 5000pts | 488: (35) | 10 |
| 6a-e, SV9 | human 3D cerebral organoids | GCamp6s | 871×871×119 | 95.8×95.8×23.8 | 28ms 800pts | 488: (428) | 30 |
| 6f-j, SV10 | *Drosophila* larval Cho neurons | jGCaMP7s | 715×715×119 | 78.7×78.7×23.8 | 5ms 500pts | 488: (428) | 100 |
| ED9, SV5 | *C. elegans* | EGFP | 793×793×119 | 87.2×87.2×23.8 | 50ms 100pts | 488: (35) | 16 |
| ED11, SV8 | Cardiac muscle cell | Fluo-8 | 949×949×119 | 104.4×104.4×23.8 | 5ms 50pts | 488: (35) | 100 |
| ED8a | Fixed zebrafish gastrula membrane | EGFP | 1001×1001×119 | 110.1×110.1×23.8 | 1.8ms 335pts | 488: (28) | 0.55 |
